# Oxidized LDL but not angiotensin II induces the interaction between LOX-1 and AT1 receptors

**DOI:** 10.1101/2020.12.14.422633

**Authors:** Li Lin, Ning Zhou, Le Kang, Qi Wang, Jian Wu, Xiaoyan Wang, Chunjie Yang, Guoping Zhang, Yunqin Chen, Hong Jiang, Ruizhen Chen, Xiangdong Yang, Aijun Sun, Hui Gong, Jun Ren, Hiroshi Aikawa, Komuro Issei, Junbo Ge, Cheng Yang, Yunzeng Zou

## Abstract

Oxidized low-density lipoprotein (Ox-LDL) can induce cardiac hypertrophy, but the mechanism is still unclear. Here we elucidate the role of angiotensin II (AngII) receptor (AT1-R) in Ox-LDL-induced cardiomycyte hypertrophy. Inhibition of Ox-LDL receptor LOX-1 and AT1-R rather than AngII abolished Ox-LDL-induced hypertrophic responses. Similar results were obtained from the heart of mice lacking endogenous Ang II and their cardiomyocytes. Ox-LDL but not AngII induced binding of LOX-1 to AT1-R, and the inhibition of LOX-1 or AT1-R rather than AngII abolished the association of these two receptors. Ox-LDL-induced ERKs phosphorylation in LOX-1 and AT1-R-overexpression cells and the binding of both receptors were suppressed by the mutants of LOX-1 (Lys266Ala/Lys267Ala) or AT1-R (Glu257Ala), however, the AT1-R mutant lacking Gq protein-coupling ability only abolished the ERKs phosphorylation. The phosphorylation of ERKs induced by Ox-LDL in LOX-1 and AT1-R-overexpression cells was abrogated by Gq protein inhibitor but not by Jak2, Rac1 and RhoA inhibitors. Therefore, the direct interaction between LOX-1 and AT1-R and the downstream Gq protein activation are important mechanisms for Ox-LDL-but not AngII-induced cardiomyocyte hypertrophy.

## Introduction

Cardiac hypertrophy is not only an adaptational state before cardiac failure, but also an independent risk factor of major cardiac events [1]. It is thus very important to understand the underlying mechanism that leads to the development of cardiac hypertrophy. Oxidized low-density lipoprotein (Ox-LDL) and lectin-like Ox-LDL receptor-1 (LOX-1), a major receptor for Ox-LDL, play fundamental roles not only in the development of atherosclerosis and hypertension [2–7], but also in the development of cardiac injuries including cardiac remodeling and heart failure [8–10].

Although LOX-1 expression has been initially identified in endothelial cells, a variety of studies have shown that it is expressed in many other cell types including cardiomyocytes. It has been reported that LOX-1 may be involved in cardiac injuries including myocardial ischemia, cardiac hypertrophy and heart failure [8–10]. LOX-1 can be upregulated and activated during myocardial ischemia, and inhibition of LOX-1 can attenuate myocardial ischemia and cardiac remodeling. LOX-1 inhibition induces reduction of oxidative stresses and inflammation responses following ischemia and thereby protects myocardium against the ischemic injury. Mechanistic studies show that inhibition of LOX-1 suppresses release of inflammatory factors and expression of angiotensin II (AngII) type 1 receptor (AT_1_-R) via inhibition of redox-sensitive pathways, leading to the improvement of cardiomyocyte hypertrophy and collagen accumulation in the ischemic regions. On the other hand, it has been indicated that there is an intimate relationship between Ox-LDL/LOX-1 and renin-angiotensin systems (RAS): Ox-LDL increases AngII converting enzyme (ACE) via LOX-1, resulting in an increase in AngII; AngII increases expression of LOX-1 through activation of AT_1_-R [11]. These reports strongly suggest that Ox-LDL/LOX-1 and RAS may cooperate in resulting in cardiac diseases including cardiac hypertrophy and heart failure. However, how Ox-LDL/LOX-1 and AngII/AT_1_-R crosstalk in the development of cardiac hypertrophy, especially how LOX1 and AT_1_-R interact during Ox-LDL-induced cardiac hypertrophy is still unclear. We therefore conduct here a study to examine the roles of AngII/AT_1_-R in Ox-LDL-induced hypertrophic responses expecially the phosphorylation of extracellular signal regulated kinases (ERKs).

## Materials and Methods

### Animal models

C57BL/6 mice were purchased from Shanghai Animal Administration Center (Shanghai, China). Ox-LDL (250 ng/kg/min), AngII (200 ng/kg/min, Sigma-Aldrich), Losartan (AT_1_-R blocker) (3 mg/kg/day, Sigma-Aldrich), LOX-1 neutralizing antibody (0.6 mg/kg/day, R&D Systems) and Enalapril (ACEI, to reduce AngII) (10 mg/kg/day, Sigma-Aldrich) were continuously administered by Alzet osmotic minipumps (DURECT) implanted subcutaneously into the mice [12, 13]. All protocols were approved by the Animal Care and Use Committee of Fudan University and in compliance with “Guidelines for the Care and Use of Laboratory Animals” published by the National Academy Press (NIH Publication No. 85-23, revised 1996).

### Lipoprotein preparation and modification of native LDL

Fresh human plasma, added citrate buffer solution containing 50 IU/ml heparin (pH 5.04), was subject to serial ultracentrifugation. After centrifugation, LDL was collected and dialyzed against phosphate buffered saline (PBS) buffer containing 200 μM EDTA for 48 hrs. Ox-LDL was prepared by incubation of LDL with 10 μM CuSO_4_ for 24 hrs at 37°C, followed by addition of 100 μM EDTA for 24 hrs, and then dialyzed in PBS buffer for 24 hrs at 4°C, changing the buffer every 8 hrs [14]. The concentration of protein was determined by the BCA protein assay kit. Oxidative modification of LDL by CuSO4 was assessed by the electrophoretic mobility of native and Ox-LDL measured on 1% agarose gel [15]. The lipoprotein pattern was visualized by staining the gel with a lipid-specific stain, and the uptake of Ox-LDL was confirmed in cultured cardiomyocytes (Supplemental Figure 1A).

### Delivery of shRNA in mice

Recombinant adeno-associated virus 9 (AAV9) vector is a kind gift from Prof. Daowen Wang at Tongji Hospital, Wuhan, China [16]. rAAV9-green fluorescent protein (GFP) (as a control), rAAV9-shLOX-1 and rAAV9-shAT_1_-R (rAAV9 generating shRNA to silence LOX-1 and AT_1_-R, respectively) were produced by a 2-plasmid protocol described previously [17] and then prepared and purified according to a published protocol [18]. The siRNA sequences were: siRNA of *LOX-1* (5’-GCAUCUCAAGUUACAGUAATT-3’; 5’-UUACUGUAACUUGAGAUGCTT-3’) and *AT1-R* (5’-GCAAAGCUGUCUUACAUUATT-3’; 5’-UAAUGUAAGACAGCUUUGCTT-3’) and their respective scramble sequences were synthesized by PCR based method and purchases from Genepharma, Inc. (Shanghai, China). The plasmid constructs were verified by sequencing. Mice were injected with rAAV-GFP, rAAV-LOX-1 or -AT_1_-R shRNA (*n* = 6 per group) through a sublingual vein injection 2 weeks before Ox-LDL infusion.

### Echocardiography and blood pressure measurements

Transthoratic echocardiography was performed using 30 MHz high frequency scanhead (VisualSonics Vevo770, VisualSonics Inc. Toronto, Canada) [12, 13]. Blood pressure (BP) was measured by a micronanometer catheter (Millar 1.4F, SPR 835, Millar Instruments, Inc.) as previously described [12, 13, 19] (See Supplementary Material for more detailed information).

### Morphology and histological analyses

Mice hearts were fixed, embedded, sectioned at 4 μm thickness and stained with hematoxylin and eosin (H-E) or relative antibodies. Cross-section area (CSA) of cardiomyocytes was measured in 20 different randomly chosen points from each cross section of left ventricle (LV) free wall (See Supplementary Material for more detailed information).

### Cell Culture, treatment, siRNA, plasmids and transfection

Neonatal cardiomyocytes of mice and COS7 cells were cultured in Dulbecco’s modified Eagle’s medium as described previously [12]. After pretreated with Losartan (10^−6^ mol/L), Enalapril (10^−6^ mol/L) or the LOX-1 neutralizing antibody (10 μg/ml) for 30 min, Ox-LDL (50 μg/ml) or vehicle was added to the cells. siRNA of *LOX-1* and AR_1_-R were synthesized by PCR based method and purchases from Genepharma, Inc. (Shanghai, China). *siRNA* was bound to siPORT Amine Transfection Reagent (Ambion) and added to culture cells (0.66 μg/ml) according to the instructions of the manufacturer. Expression vectors encoding wild type AT_1_-R and its K199Q (replacement of Lys199 with Glu), Q257A (replacement of Glu257 with Ala), C289A (replacement of Cys 289 with Ala) and AT1-Ri2m (lacks a binding domain for G protein) mutants were constructed as described previously [12, 20, 21]. The expression vector encoding LOX-1 was a gift from Dr. Tatsuya Sawamura at National Cardiovascular Center Research Institute, Osaka, Japan [22]. Dominant negaive mutants of LOX-1 (DN-LOX-1: Lys266 and Lys267 in LOX-1 were substituted by alanine) were constructed as described in a previous study [23]. The mutated genes were sequenced completely to verify the mutations. Transfection of plasmids was performed by using lipofectamine 2000™ transfection reagent (Invitrogen) according to the manufacturer’s instructions.

### Real-time RT-PCR

Purified RNA was subjected to a real-time RT-PCR analysis. A comparative CT method was used to determine relative quantification of RNA expression [24]. All PCR reactions were performed at least triplicate.

### Western blot analyses

Total proteins extracted from the hearts or cells were subjected to Western blot analysis for phosphorylation of ERKs, total ERKs, β-Actin or GAPDH. The amounts of LOX-1 and AT_1_-R were examined after dividing the membrane fraction and the cytosolic fraction. Following antibodies were used to detect protein expression: anti phosphorylated ERKs (Cell Signaling Technology), anti total ERKs (Santa Cruz Biotechnology), anti LOX-1 (Abcam), anti AT_1_-R (Santa Cruz Biotechnology).

### Immunoprecipitation

Proteins of cellular membrane were extracted using the ProteoExtract^®^ Transmembrane Protein Extraction Kit (NOVAGEN). Total 500 μg of membrane proteins were used for immunoprecipitation with an anti AT_1_-R or LOX-1 antibody. The immunecomplexes were precipitated with protein A/G PLUS-Agrose (Santa Cruz Biotechnology) and subjected to the SDS-PAGE for detecting amounts of LOX-1 or AT_1_-R.

### Immunofluorescence

Cardiomyocytes cultured on glass cover slides in serum-free DMEM for 24 hrs were incubated with anti-MHC (Upstate), anti-LOX-1 (Santa Cruz Biotechnology) and anti-AT_1_-R (Abcam) antibodies, and then with secondary antibodies conjugated with FITC or Alex (Invitrogen) according to the manufacturer’s directions. The surface area of cardiomyocytes stained by MHC was determined with image analysis software (Leica Qwin 3) and calculated as the mean of 100 to 120 cells from randomly selected fields.

### Fusion protein design and bimolecular fluorescence complementation (BiFC) assay

The following expression fusion plasmids were constructed for this assay: AT_1_/KGC, AT_1_-R tagged by subunit C of GFP at C terminals of the receptor; AT_1_/KGN, AT_1_-R tagged by subunit N of GFP at C terminals of the receptor; LOX-1/KGN, LOX-1 receptor tagged by subunit N of GFP at N terminals of the receptor; LOX-1/KGC, LOX-1 receptor tagged by subunit C of GFP at N terminals of the receptor. These plasmids were constructed as previously described [25] using the CoraHue® FLUO-chase Kit (Medical & Biological Laboratories Co., LTD., Japan). AT_1_/KGC and LOX-1/KGN or AT_1_/KGN and LOX-1/KGC were co-transfected into the COS7 cells.

### Statistical analyses

All statistical significance was determined by one-way or two-way ANOVA with Newman-Keuls test for post-hoc analysis. A *p* value of < 0.05 was considered statistically significant. All results are presented as mean ± standard error of the mean (SEM).

## Results

### Ox-LDL-induced cardiomyocyte hypertrophy is inhibited by a LOX-1 or an AT1-R blocker but not by reduction of AngII

We first confirmed the induction of cardiomyocyte hypertrophy by Ox-LDL. Subcutaneous infusion of Ox-LDL for 2 weeks, which resulted in a significant elevation of plasma Ox-LDL (Supplemental Figure 2A), induced a definite cardiac hypertrophy in male C57BL/6 mice, such as the increases in LV wall thickness, gross heart size, heart weight to body weight (HW/BW) ratio, cross-sectional area (CSA) of cardiomyocytes evaluated by echocardiography and histology (Figure 1A, B), although it did neither elevate BP nor result in atherosclerosis in these mice (Supplemental Figure 2B, C). Incubation of cultured cardiomyocytes with Ox-LDL also induced significant increases in cardiomyocyte size evaluated by surface area of α-myosin heavy chain (α-MHC)-stained cells (Figure 1E). Phosphorylation of protein kinases such as ERKs and reprogramming of expression of some fetal genes such as *atrial natriuretic peptide* (*ANP*) and *skeletal α-actin* (*SAA*) are also important hypertrophic responses in cardiomyocytes [19, 26, 27]. Either infusion of Ox-LDL into mice or addition of Ox-LDL to cultured cardiomyocytes induced upregulation of both the phosphorylation of ERKs and the expression of *ANP* and *SAA* genes (Figure 1C, D, F, G), respectively. These results collectively confirmed the induction of cardiomyocyte hypertrophy by Ox-LDL.

**Figure 1.**
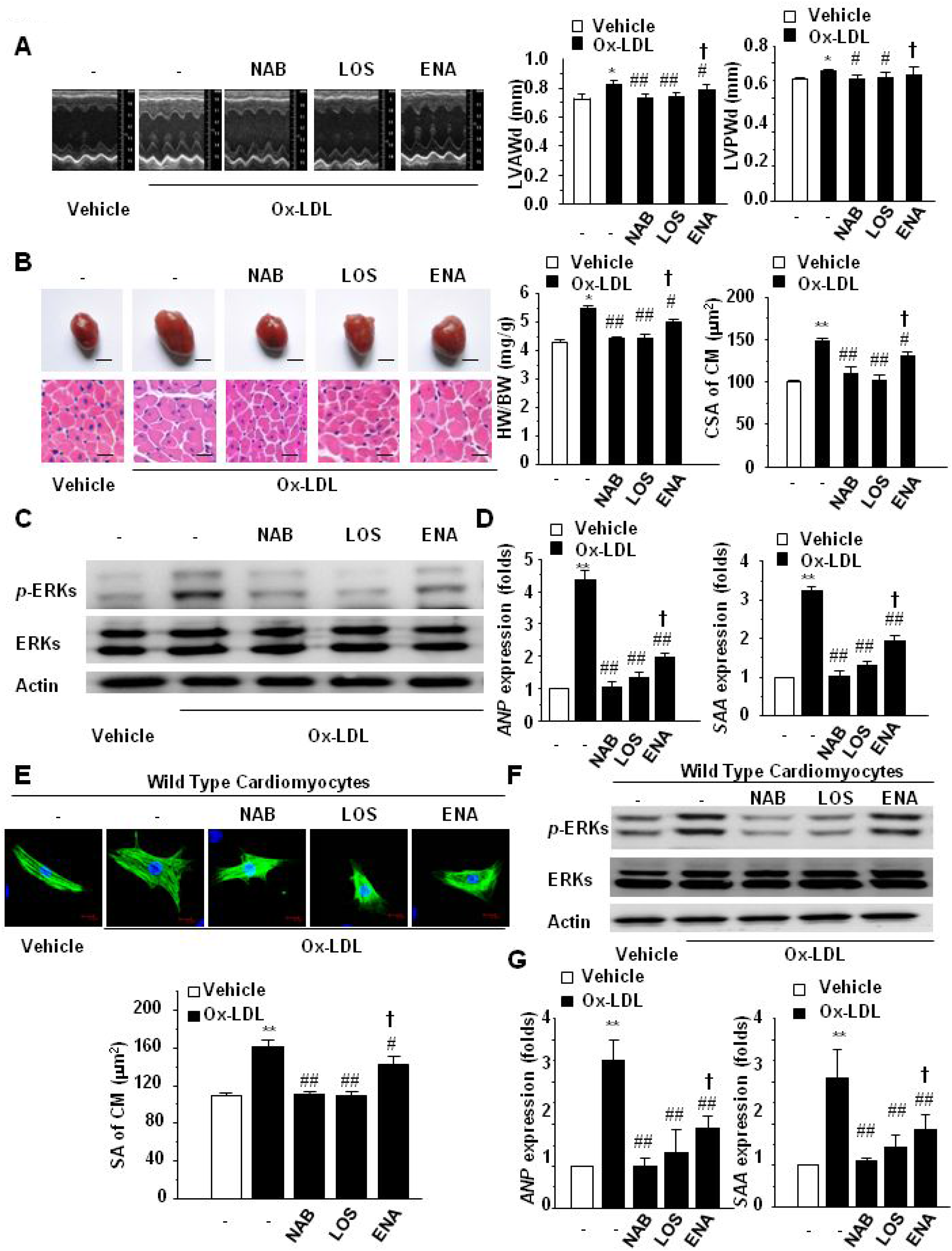
Involvement of LOX-1, AT_1_-R and AngII in Ox-LDL-induced cardiomyocyte hypertrophy. **A**-**D**, Adult male C57BL/6 mice pretreated with a neutralizing antibody of LOX-1 (NAB), Losartan (LOS), Enalapril (ENA) or vehicle (−), were infused with vehicle or Ox-LDL for 2 weeks. **A**, Echocardiograghic analysis. Representative M-mode tracings are shown. LVAWd, left ventricular (LV) anterior wall thickness at end-diastole; LVPWd, LV posterior wall thickness at end-diastole. **B**, Heart morphology and H-E stained LV sections. Representative global heart photographs (scale bar: 2 mm) and H-E stained heart sections (scale bar: 20 μm) are shown. Heart weight to body weight (HW/BW) was measured from 5 mice. Cross sectional area (CSA) of cardiomyocytes was measured from 5 sections for one heart. **C**, Protein kinase phosphorylation after infusion with Ox-LDL. Phosphorylation of ERKs in the heart of mice was examined by Western blotting using an anti-phosphor-ERKs antibody. Total ERKs and Actin were used as loading controls. Representative photograms are shown. **D**, Specific gene expression. mRNA expression of *ANP* and *SAA* in heart tissues of mice were evaluated by the real time RT-PCR method. All data are expressed as mean ± SEM of 5 mice (n=5) * *p* < 0.05, ** *p* < 0.01 vs. mice with vehicle; # *p* < 0.05, ## *p* < 0.01 vs. mice without pretreatment (−) but with Ox-LDL infusion. † *p* < 0.05 vs. mice with vehicle. **E**-**G**, Cultured cardiomyocytes (CM) of neonatal mice, pretreated without (−) or with the NAB, LOS or ENA for 30 min, were exposed to vehicle or Ox-LDL for 24~48 hrs. **E**, CM morphology ang size. Representative immunofluorescent staining for α-MHC (green staining) and DAPI staining (blue) are shown (scale bar: 10 μm). Surface area (SA) of CM was evaluated by measuring 100 CM in one dish. **F**, Protein kinase activation and **G**, Specific gene expression after Ox-LDL stimulation. Phosphorylation of ERKs was examined by Western blotting using an anti-phosphor-ERKs antibody. Total ERKs and Actin were used as loading controls. Representative photograms are shown. mRNA expression of *ANP* and *SAA* were evaluated by the real time RT-PCR method. All data are expressed as mean ± SEM of 5 independent experiments (n=5). ** *p* < 0.01 vs. CM with vehicle; # *p* < 0.05, ## *p* < 0.01 vs. CM without pretreatment (−) but with Ox-LDL stimulation. † *p* < 0.05 vs. CM with vehicle.

We next examined the role of Ox-LDL receptor LOX-1 in Ox-LDL-induced cardiac hypertrophy using a LOX-1 neutralizing antibody (NAB) to block the effect of LOX-1. Administration of the LOX-1 neutralizing antibody in mice resulted in a great inhibition of cardiac hypertrophic responses induced by Ox-LDL (Figure 1A, B). Phosphorylation of ERKs and gene expression of *ANP* and *SAA* were also downregulated by the LOX-1 neutralizing antibody in hearts of mice (Figure 1C, D), although the plasma Ox-LDL levels and the BP in these mice were not affected by the neutralizing antibody (Supplemental Figure 2A, B). Moreover, in cultured cardiomyocytes, pretreatment with the LOX-1 neutralizing antibody also significantly abolished ox-LDL-induced increases in cell size, ERKs phosphorylation and mRNA expression of *ANP* and *SAA* (Figure 1E-G). All these results collectively suggest that Ox-LDL induces cardiomyocyte hypertrophy through LOX-1.

We further examined the roles of AngII and AT_1_-R in Ox-LDL-induced cardiac hypertrophy using an AT_1_-R blocker, Losartan (LOS) and an ACE inhibitor, Enalapril (ENA), respectively. We first confirmed the effects of Ox-LDL on AngII expression in mice and cultured cardiomyocytes. Ox-LDL obviously increased AngII levels in the plasma of mice, which could be suppressed by LOX-1 neutralizing antibody and Enalapril but not by Losartan (Supplemental Figure 3A). Interestingly, Ox-LDL only induced a marginal increase in AngII in the supernant of cardiomyocytes (Supplemental Figure 3B). Both *in vivo* and *in vitro* experiments showed that the development of cardiomyocyte hypertrophy induced by Ox-LDL was significantly abrogated by Losartan, however, Enalapril only slightly attenuated Ox-LDL-induced cardiomyocyte hypertrophy (Figure 1A-G), although the both drugs did not affect the plasma Ox-LDL levels and the BP in Ox-LDL-treated mice (Supplemental Figure 2A, B) and could similarly depress the BP in hypertensive mice (Supplemental Figure 3C). These data suggest that AT_1_-R rather than AngII is involved in Ox-LDL-induced cardiomyocye hypertrophy, stimulating our interest on exploring the mechanism for these results.

### Ox-LDL induces hypertrophic responses through LOX-1 and AT1-R under the condition lacking AngII

To investigate the mechanism by which Ox-LDL induces cardiac hypertrophic responses through LOX-1 and AT_1_-R, we first confirmed the effects of Ox-LDL, LOX-1 and AT_1_-R under the condition without the involvement of AngII. We therefore used *angiotensinogen* gene (*ATG*) knockout mice lacking endogenous AngII [12]. Ox-LDL could induce both *in vivo* and *in vitro* cardiomyocyte hypertrophic responses in the *ATG* knockout mice, and all these hypertrophic responses were significantly suppressed by either the LOX-1 neutralizing antibody or the AT_1_-R blocker (Figure 2A-G). We also used siRNA method to downregulate LOX-1 and AT_1_-R expressions both in mouse heart and cultured cardiomyocytes (Supplemental Figure 4A, B). Consistent with the results of pharmacological inhibitors, both *in vivo* and *vitro* experiments showed that knocking down of either LOX-1 or AT_1_-R could significantly inhibit all hypertrophic responses induced by Ox-LDL (Figure 3A-F). These data strongly confirmed that Ox-LDL can induce cardiac hypertrophy through LOX-1 and AT_1_-R independently of AngII. Previous studies have indicated that Ox-LDL upregulates the expression of ACE through LOX-1 pathway, which increases local AngII and then activates AT_1_-R [28–30]. We here speculate a novel mechanism by which Ox-LDL induces hypertrophic responses through AT_1_-R rather than AngII.

**Figure 2.**
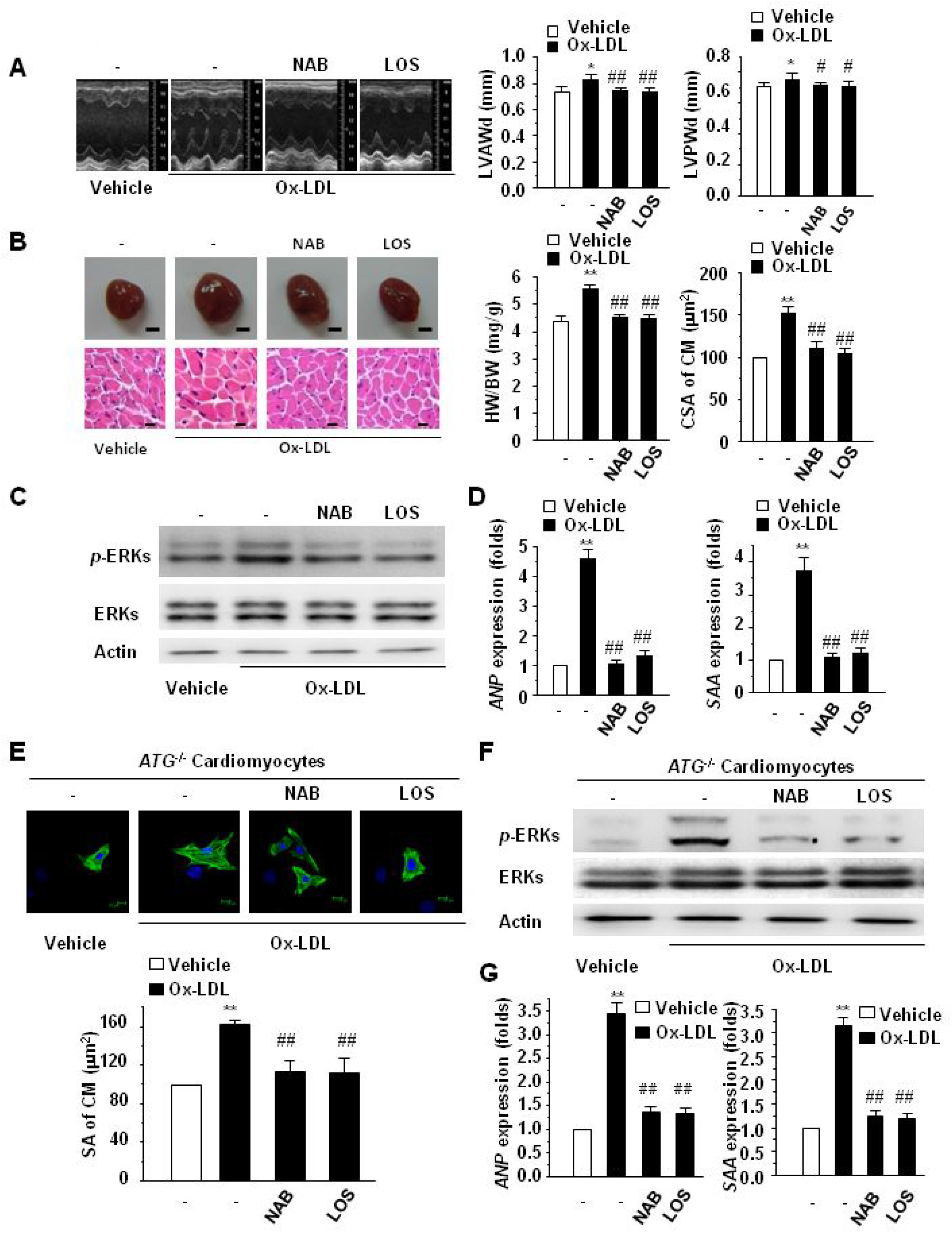
Induction of cardiomyocyte hypertrophy by Ox-LDL in *ATG*^−/−^ mice**. A**-**D**, Adult male *ATG*^−/−^ mice pretreated with the NAB, LOS or vehicle (−) were infused with vehicle or Ox-LDL for 2 weeks. **A**, Echocardiograghic analysis. Representative M-mode tracings are shown. **B**, Heart morphology and H-E stained LV sections. Representative global heart photographs (scale bar: 2 mm) and H-E stained heart sections (scale bar: 20 μm) are shown. HW/BW was measured from 5 mice. CSA of cardiomyocytes was measured from 5 sections for one heart. **C**, Protein kinase phosphorylation after infusion with Ox-LDL. Phosphorylation of ERKs in the heart of mice was examined by Western blotting using an anti-phosphor-ERKs antibody. Total ERKs and Actin were used as loading controls. Representative photograms are shown. **D**, Specific gene expression. mRNA expression of *ANP* and *SAA* in heart tissues of mice were evaluated by the real time RT-PCR method. All data are expressed as mean ± SEM of 5 mice (n=5). * *p* < 0.05, ** *p* < 0.01 vs. mice with vehicle; # *p* < 0.05, ## *p* < 0.01 vs. mice without pretreatment (−) but with Ox-LDL infusion. **E-G**, Cultured cardiomyocytes (CM) of neonatal *ATG*^−/−^ mice, pretreated without (−) or with the NAB or LOS for 30 min, were exposed to vehicle or Ox-LDL for 24~48 hrs. **E**, CM morphology and size. Immunofluorescent staining for α-MHC (green staining) and DAPI staining (blue) are shown (scale bar: 10 μm). Surface area (SA) of CM was evaluated by measuring 100 CM in one dish. **F**, Protein kinase activation and **G**, specific gene expression after Ox-LDL stimulation. Phosphorylation of ERKs was examined by Western blotting using an anti-phosphor-ERKs antibody. Total ERKs and Actin were used as loading controls. Representative photograms are shown. mRNA expression of *ANP* and *SAA* were evaluated by the real time RT-PCR method. All data are expressed as mean ± SEM of 5 independent experiments (n=5). ** *p* < 0.01 vs. CM with vehicle; # *p* < 0.05, ## *p* < 0.01 vs. CM without pretreatment (−) but with Ox-LDL stimulation.

**Figure 3.**
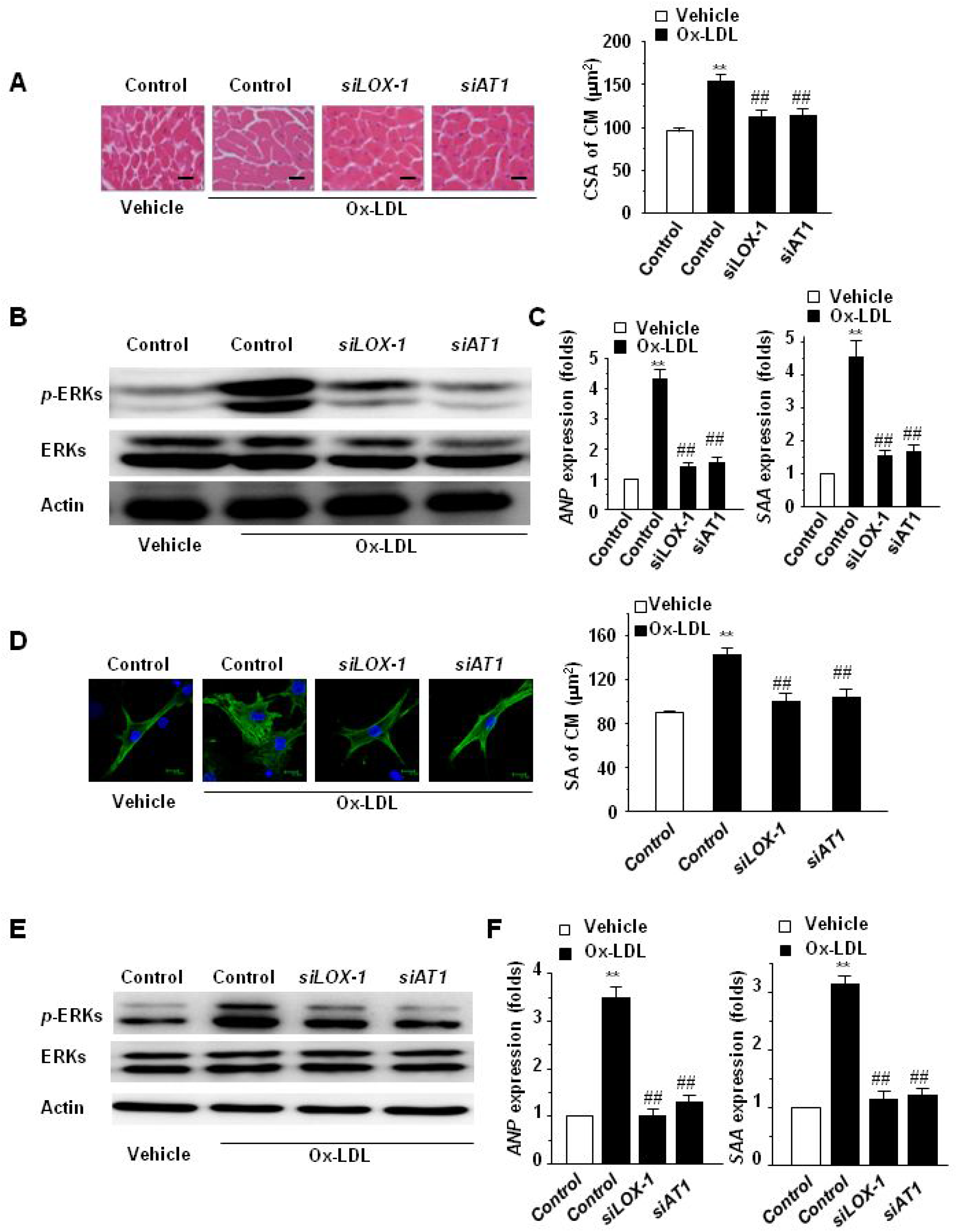
Effects of knockdown of LOX-1 and AT_1_-R on Ox-LDL-induced cardiomyocyte hypertrophy in *ATG*^−/−^ mice. **A**-**C**, Adult male *ATG*^−/−^ mice pretreated with AAV9-mediated shRNA of LOX-1 (*siLOX-1*) or AT_1_-R (*siAT_1_-R*) or control were infused with vehicle or Ox-LDL for 2 weeks. **A**, H-E stained LV sections. Representative H-E stained heart sections (scale bar: 20 μm) are shown. CSA of cardiomyocytes was measured from 5 sections for one heart. **B**, Protein kinase phosphorylation. Phosphorylation of ERKs in the heart of mice was examined by Western blotting using an anti-phosphor-ERKs antibody. Total ERKs and Actin were used as loading controls. Representative photograms are shown. **C**, Specific gene expression. mRNA expression of *ANP* and *SAA* in heart tissues of mice were evaluated by the real time RT-PCR method. All data are expressed as mean ± SEM of 5 mice (n=5). ** *p* < 0.01 vs. mice with control and vehicle; ## *p* < 0.01 vs. mice with control siRNA and Ox-LDL treatment. **D**-**F**, Cultured cardiomyocytes (CM) of adult male *ATG*^−/−^ mice, added by siRNA of LOX-1 (*siLOX-1*), AT_1_-R (*siAT_1_-R*) or control for 24 hrs, were exposed to vehicle or Ox-LDL for 24~48 hrs. **D**, CM morphology and size. Immunofluorescent staining for α-MHC (green staining) and DAPI staining (blue) are shown (scale bar: 10 μm); Surface area (SA) of CM was evaluated by measuring 100 CM in one dish. **E**, Protein kinase phosphorylation and **F**, specific gene expression. Phosphorylation of ERKs was examined by Western blotting using an anti-phosphor-ERKs antibody. Total ERKs and Actin were used as loading controls. Representative photograms are shown; mRNA expression of *ANP* and *SAA* were evaluated by the real time RT-PCR method. All data are expressed as mean ± SEM of 5 independent experiments (n=5). ** *p* < 0.01 vs. CM with control and vehicle; ## *p* < 0.01 vs. CM without siRNA (Control) but with Ox-LDL stimulation.

### Syntropic upregulation and co-localization of LOX-1 and AT1-R induced by Ox-LDL

We next asked how LOX-1 and AT_1_-R are involved in Ox-LDL-induced cardiomyocyte hypertrophy. The expressions of LOX-1 and AT_1_-R proteins were both upregulated not only in hearts of mice infused with Ox-LDL but also in cultured cardiomyocytes incubated with Ox-LDL (Figure 4A-C). Inhibition of either LOX-1 or AT_1_-R alone by LOX-1 neutralizing antibody or Losartan, respectively, could significantly suppress Ox-LDL-induced upregulation of both the receptors, whereas inhibition of ACE by Enalapril did not exert similar effects (Figure 4A-C). The upregulation of LOX1 and AT_1_-R by Ox-LDL and the inhibitory effects of LOX-1 neutralizing antibody and Losartan on the upregulation were also observed in the heart and cultured cardiomyocytes of *ATG* knockout mice (Figure 4D, E). In agreement with the results obtained from the pharmacological inhibitors, knocking down of either LOX-1 or AT_1_-R alone could significantly abrogate Ox-LDL-induced upregulation of the two receptors (Figure 4F, G). Moreover, immunofluorescent staining of LOX-1 (Red) and AT_1_-R (Green) revealed that the signals of LOX-1 as well as AT_1_-R were stronger in cardiomyocytes incubated with Ox-LDL than in those without Ox-LDL incubation, both of which displayed more co-localization nearly the cellular membrane (Figure 4C). Either the strength of the signals or the co-localization of the two receptors in Ox-LDL-stimulated cardiomyocytes became significantly less when pretreated with the LOX-1 neutralizing antibody or Losartan, but Enalapril pretreatment could not reduce the Ox-LDL-stimulated increases in the expression and the co-localization of the two receptors (Figure 4C).

**Figure 4.**
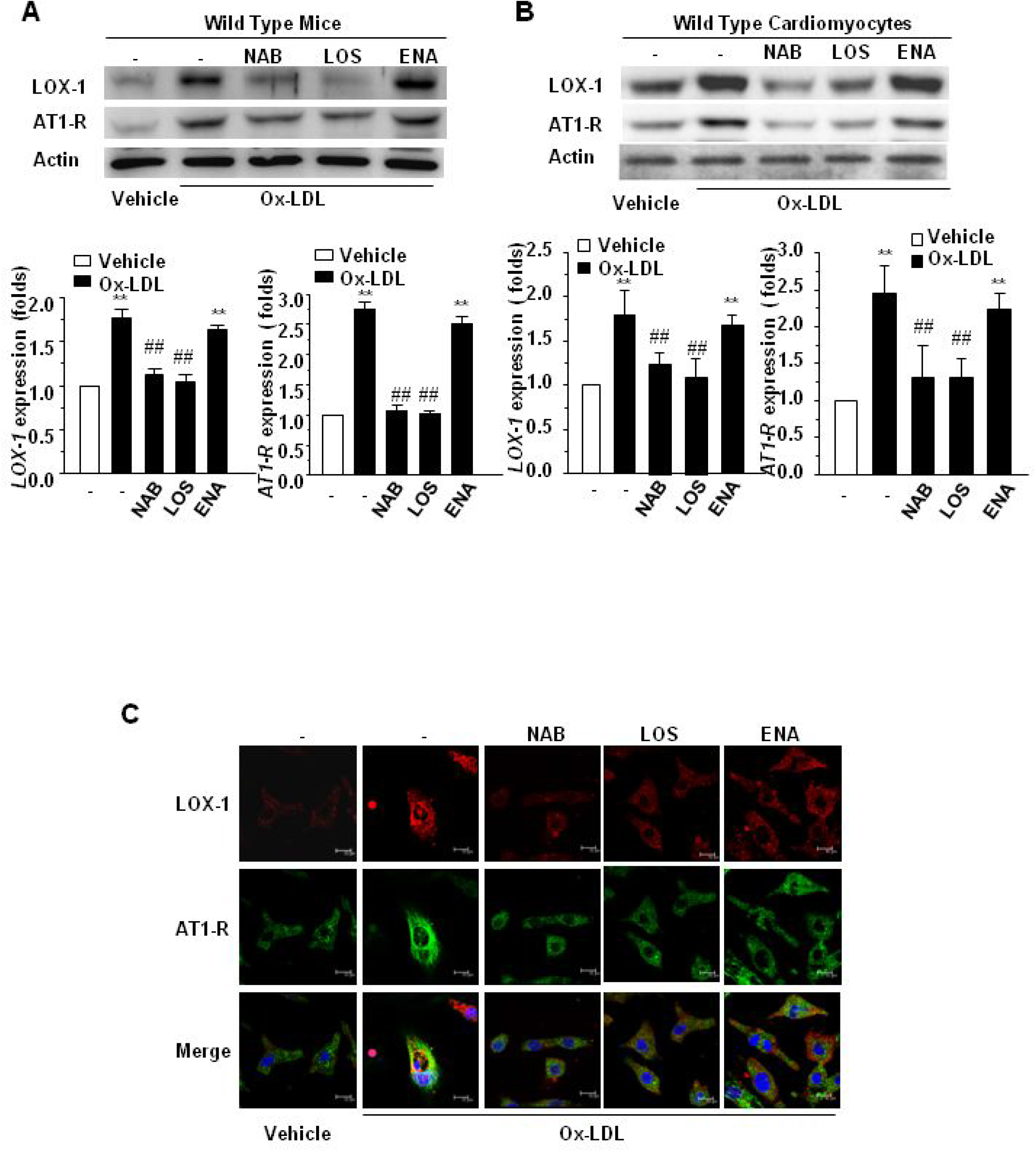
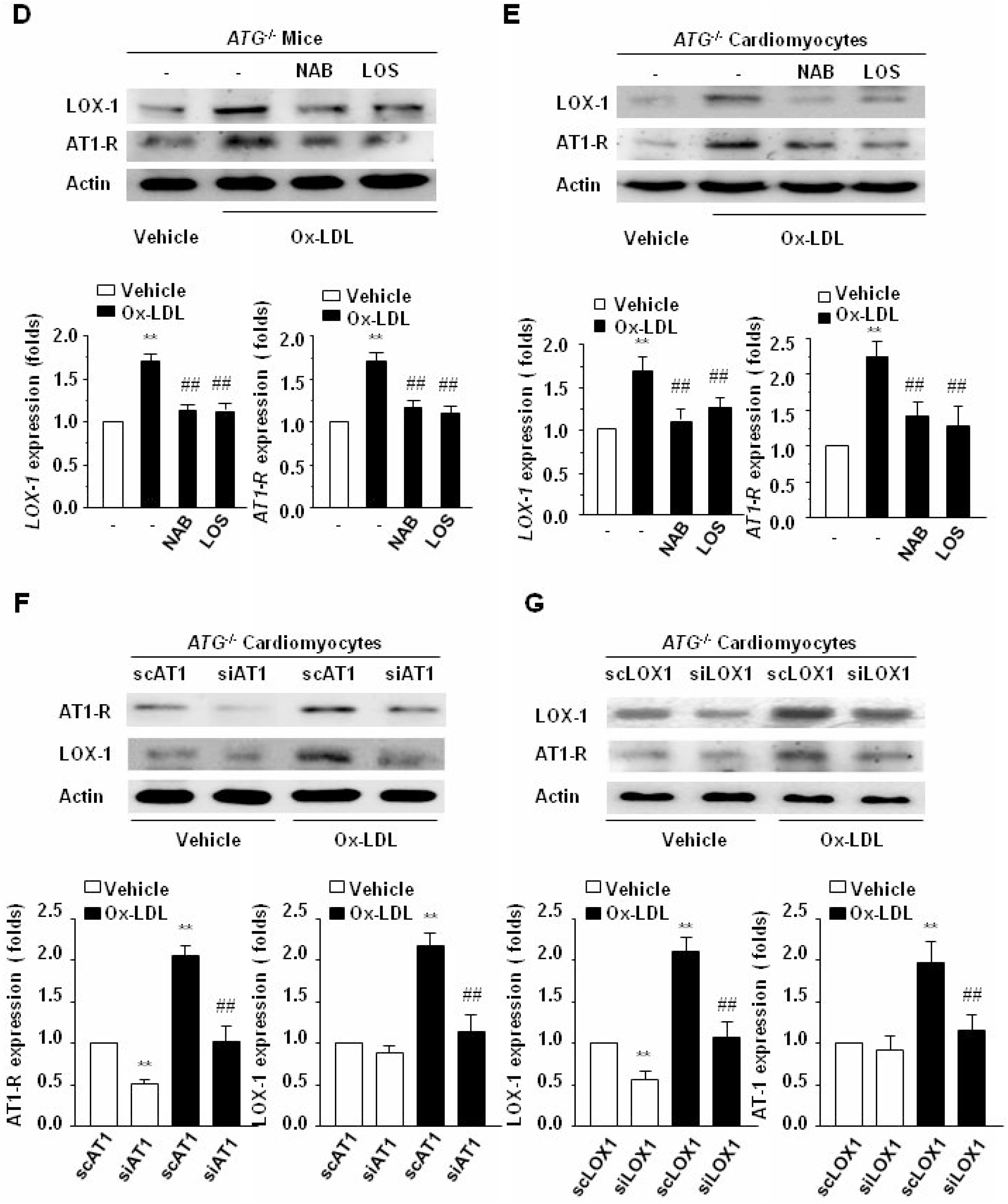
Expression and co-localization of LOX-1 and AT_1_-R induced by Ox-LDL. **A**, Wild type or **D**, *ATG*^−/−^mice, pretreated with the NAB, LOS, ENA or vehicle (−) were infused with vehicle or Ox-LDL for 2 weeks. **B** and **C**, Cultured cardiomyocytes (CM) of wild type or **E** and **G**, *ATG*^−/−^ mice, pretreated with the NAB, LOS ENA or vehicle (−) for 30 min, were exposed to vehicle or Ox-LDL for 24 hrs. **A** and **B, D**-**G**, Membrane proteins were extracted from LV of mice and CM and subjected to Western blot analyses for LOX-1 and AT_1_-R protein expression using anti LOX-1 and AT_1_-R antibodies, respectively. β-actin in whole cell lysate was used as the loading control. Representative photograms from 5 hearts or 5 independent experiments are shown. **A** and **D**, mRNA isolated from the similarly treated heart tissues and **B** and **E**, CM was analyzed by real time RT-PCR for *LOX-1* and *AT*_*1*_*-R* gene expression. Data are expressed as mean ± SEM of 5 mice or 5 experiments (n=5). ** *p* < 0.01 vs. respective mice and CM with vehicle; ## *p* < 0.01 vs. respective mice and CM without pretreatment (−) but with Ox-LDL. **C**, Immunofluorescent staining for LOX-1 (red) and AT_1_-R (green) in CM. Representative photographs from 5 independent experiments are shown. Blue, DAPI staining; Scale bar, 10 μm.

### Association of LOX-1 and AT1-R induced by Ox-LDL

We therefore asked the relationship between LOX-1 and AT_1_-R after stimulation with Ox-LDL. Do they interact directly or through other mediators? To answer this question we employed the method of co-immuneprecipitation using both anti LOX-1 and anti AT_1_-R antibodies. Western blot analyses of immunecomplexes precipitated with an antibody against AT_1_-R of heart tissue (Figure 5A, top) and cultured cardiomyocytes (Figure 5B, top) for LOX-1 expression showed an enhanced association of LOX-1 with AT_1_-R after simulation with Ox-LDL. Treatment with the LOX-1 neutralizing antibody or Losartan almost completely abolished the association. Enalapril, however, had no markedly inhibitory effect on this association. Similar results were obtained in analyses of immunecomplexes precipitated with an anti LOX-1 antibody for AT_1_-R expression (Figure 5A, B, bottom). Furthermore, the increase in the association of LOX-1 with AT_1_-R after Ox-LDL stimulation and the inhibition of this increase by the LOX-1 neutralizing antibody and Losartan were also observed in the hearts (Figure 5C) and cultured cardiomyocytes (Figure 5D) of *ATG* knockout mice. Consistent with the pharmacological effects, siRNA of LOX-1 or that of AT_1_-R significantly reduced Ox-LDL-stimulated association of LOX-1 with AT_1_-R in the hearts and cultured cardiomyocytes of *ATG* knockout mice (Figure 5E, F). Additionally, to ask whether AngII could also stimulate association of AT_1_-R with LOX-1, AngII was administrated to mice and cultured cardiomyocytes. The results revealed that AngII could not induce association of AT_1_-R with LOX-1 in hearts of mice (Supplemental Figure 5, top) nor in cultured cardiomyocytes (Supplemental Figure 5, bottom). All these findings suggest that there exists an association of LOX-1 with AT_1_-R initiated by Ox-LDL stimulation but not by AngII in cardiomyocytes.

**Figure 5.**
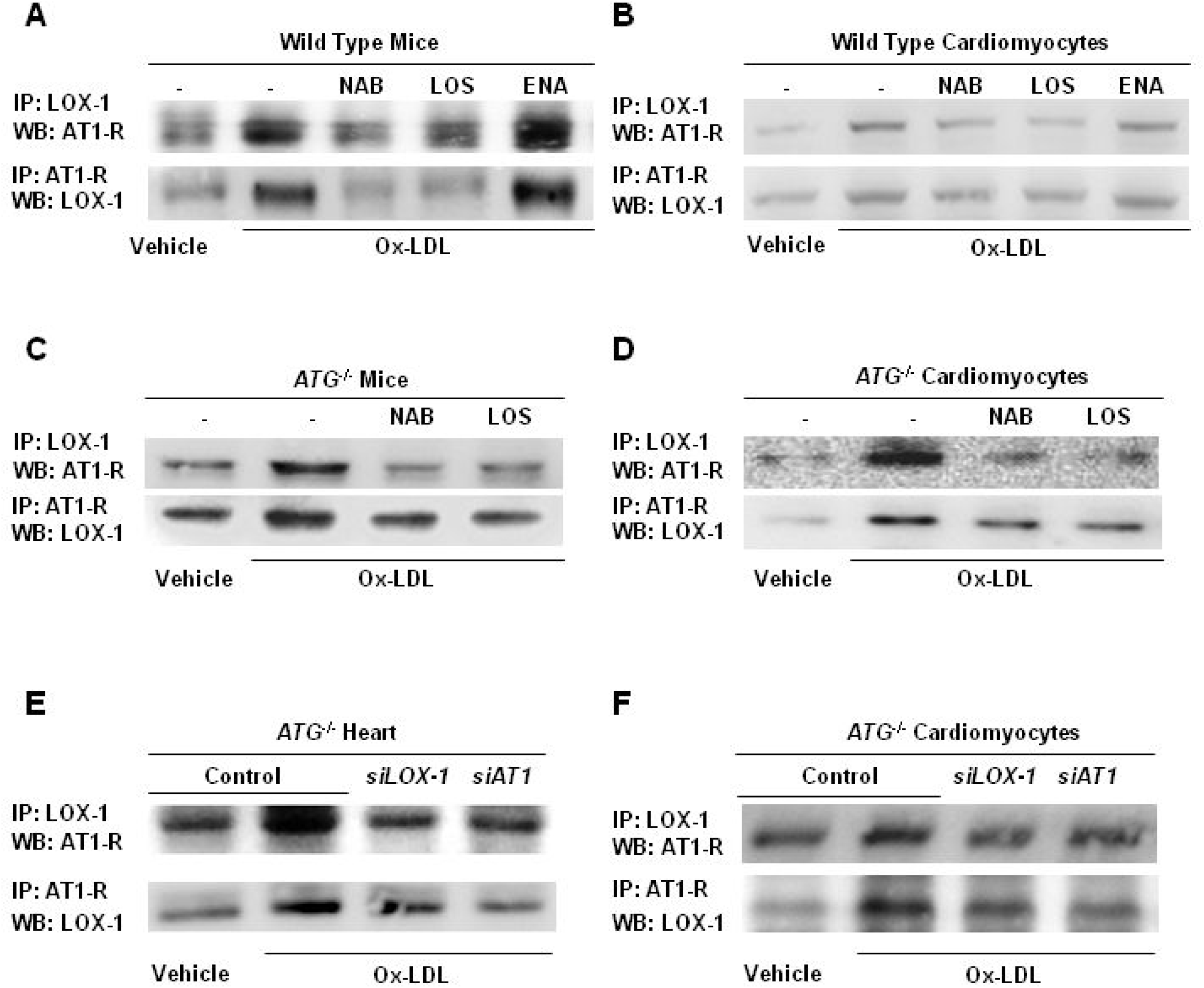
Co-immunoprecipatation of LOX-1 and AT_1_-R. **A**, Wild type or **C**, *ATG*^−/−^ mice, pretreated with the NAB, LOS, ENA or vehicle (−), were infused with vehicle or Ox-LDL for 2 weeks. **B**, Cultured cardiomyocytes (CM) of wild type or **D**, *ATG*^−/−^ mice, pretreated with the NAB, LOS, ENA or vehicle (−) for 30 min, were exposed to vehicle or Ox-LDL for 24 hrs. **E**, *ATG*^−/−^ mice or CM of *ATG*^−/−^ mice, treated by siRNA of *LOX-1* (*siLOX-1*), *AT*_*1*_*-R* (*siAT_1_-R*) or vehicle (Control), were exposed to vehicle or Ox-LDL. Membrane proteins extracted from LV of mice or CM were immunoprecipated (IP) with antibodies against LOX-1 or AT_1_-R and the immunecomplexes were subjected to Western blot (WB) analyses for AT_1_-R and LOX-1, respectively. Representative photograms from 5 hearts or 5 independent experiments are shown.

### Involvement of LOX-1, AT1-R and downstream molecules in Ox-LDL-stimulated ERKs phosphorylation

Because of a presumed relation between LOX-1 and AT_1_-R that is independent of AngII during Ox-LDL-induced cardiomyocyte hypertrophy, COS7 cells, which lack expressions of endogenous AngII, AT_1_-R [12] and LOX-1 (Supplemental Figure 6), were used to confirm the gain of function of the two receptors and the mechanism of how LOX-1 and AT_1_-R work under the stimulation with Ox-LDL. We transiently transfected the plasmids of LOX-1, AT_1_-R or both of them into cultured COS7 cells and then stimulated the cells with Ox-LDL. Ox-LDL did not induce significant increases in the phosphorylation of ERKs in COS7 cells without any transfection nor in those transfected with LOX-1 or AT_1_-R alone, whereas the phosphorylation of ERKs was largely induced by Ox-LDL in those doubly transfected with LOX-1 and AT_1_-R (Figure 6A). However, although AngII could induce increases in the phosphorylation of ERKs in COS7 cells doubly transfected with LOX-1 and AT_1_-R, the ERK phosphorylation was not significantly different from that in the cells transfected with AT_1_-R alone (Supplemental Figure 7A). Additionally, AngII could not induce ERKs phosphorylation in the COS7 cells transfected with LOX-1 alone (Supplemental Figure 7A). These results indicate that co-expression of LOX-1 with AT_1_-R provides COS7 cells with the ability to respond to Ox-LDL which is independent of AngII.

**Figure 6.**
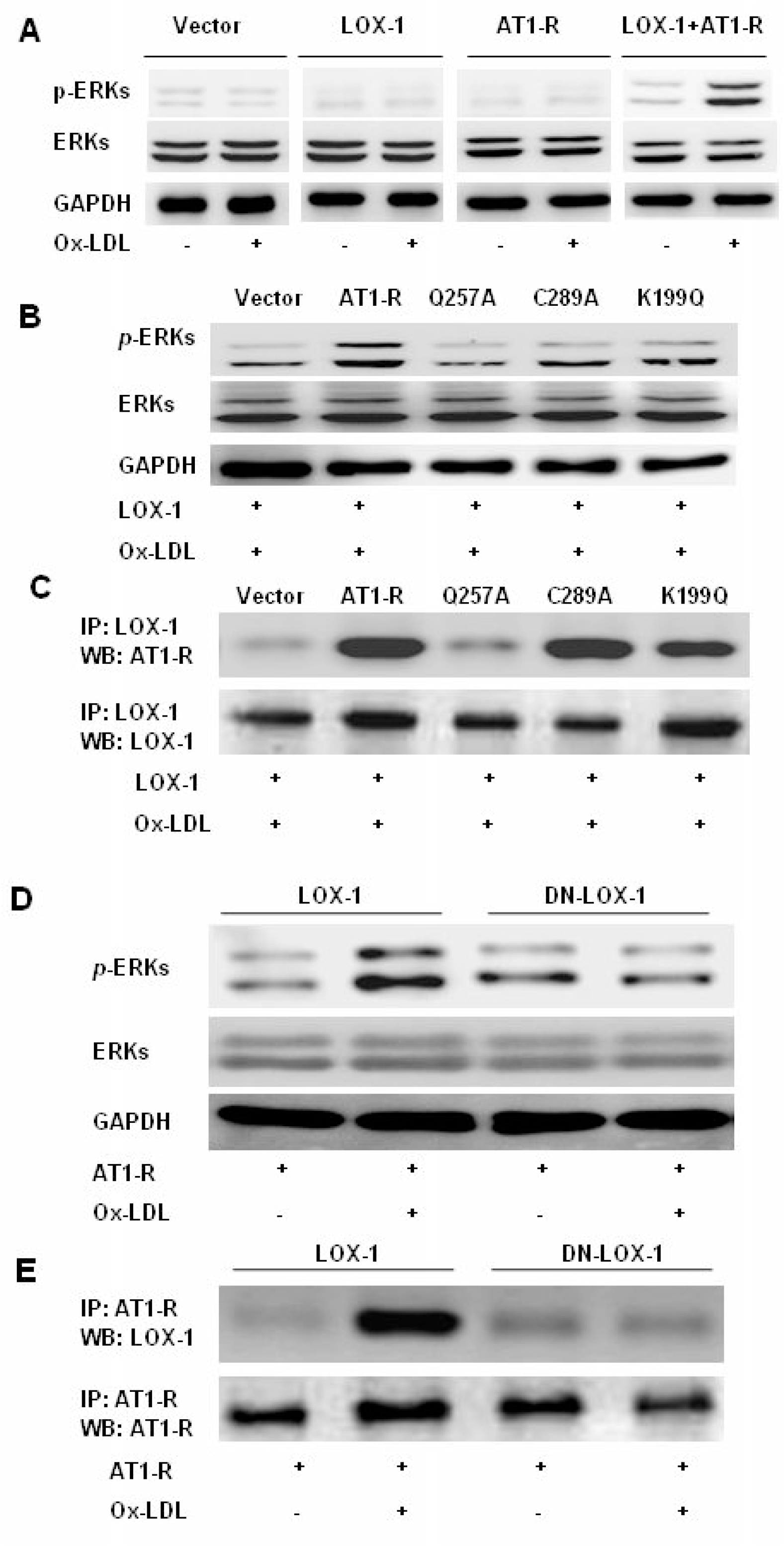
Effects of co-transfection of LOX-1, AT_1_-R or their mutants on ERKs phosphorylation and association of the two receptors. **A**, COS7 cells were transiently transfected with empty vector, *LOX-1* or *AT*_*1*_*-R* alone, or co-transfected with both *LOX-1* and *AT*_*1*_*-R* for 24 hrs and stimulated by Ox-LDL or vehicle for 10 min. **B** and **C**, COS7 cells were transiently co-transfected by *LOX-1* with *AT*_*1*_*-R* mutants K199Q, G257A or C289A and stimulated by Ox-LDL. **D** and **E**, COS7 cells were transiently co-transfected by *AT*_*1*_*-R* with wild type *LOX-1* or *DN-LOX-1* and stimulated by Ox-LDL or vehicle. Phosphorylation of ERKs was detected by an anti phosphor-ERKs antibody. Total ERKs and GAPDH were used as loading controls. Membrane proteins were immunoprecipated (IP) with antibodies against LOX-1 and the immunecomplexes were subjected to Western blot (WB) analyses for AT_1_-R and LOX-1, respectively. Representative photograms from 5 independent experiments are shown.

We next examined the mechanisms by which Ox-LDL induces ERKs phosphorylation through AT_1_-R and LOX-1 activation by using relevent mutants of the receptors. Three AT_1_-R mutants, which respectively inhibit AngII-dependent (K199Q) or -independent (Q257A, C289A) activation of the receptor and the dominant negative mutant of LOX-1(DN-LOX-1: K266A/K267A) [12, 20, 21] were constructed and used. The wild type AT_1_-R or three mutants of AT_1_-R was co-transfected into COS7 cells with LOX-1, or DN-LOX-1 was co-transfected into COS7 cells with wild type AT_1_-R, and then the cells were stimulated with Ox-LDL. Ox-LDL induced significant increases in phosphorylation of ERKs in wild type AT_1_-R-co-transfected COS7 cells, and the increases were declined in Q257A-co-transfected cells but maintained in either K199Q- or C289A-co-transfected ones (Figure 6B), however, AngII-induced activation of ERKs in AT_1_-R-co-transfected cells was inhibited only in K199Q-co-transfected cells (Supplemental Figure 7B). Consisting with the ERKs phosphorylation, association of the two receptors stimulated by Ox-LDL was disappeared in Q257A-co-transfected cells and maintained in K199Q- or C289A-co-transfected ones (Figure 6C), suggesting that Ox-LDL-induced association of AT_1_-R with LOX-1 and subsequent activation of ERKs was dependent on Q257 of AT1-R. Moreover, Ox-LDL-induced increases in association of LOX-1 with AT_1_-R and the phosphorylation of ERKs were abolished in DN-LOX-1-co-transfected COS7 cells (Figure 6D, E).

To ask which downstream molecules mediate activation of ERKs from Ox-LDL-stimulated LOX-1/AT_1_-R association, we first used inhibitors for Gq protein α subunit and Jak2, which have been shown to mediate activation of ERKs by AngII-independent activation of AT_1_-R [12], and inhibitors for Rac1 and RhoA, which are known as the downstream molecules of LOX-1 [31]. Interestingly, the Inhibitor for Gq protein α subunit but not that for Jak2, Rac1 or RhoA abrogated the Ox-LDL-induced phosphorylation of ERKs (Figure 7A). To confirm the pharmacological effect, we co-transfected LOX-1 with a AT_1_-R mutant, AT_1_-Ri2m, which lacks the ability coupling with the Gq protein. Although Ox-LDL-induced the association of LOX-1 with AT_1_-R were not affected (Figure 7C), the phosphorylation of ERKs were abolished by AT_1_-Ri2m transfection (Figure 7B). These results clearly suggest that coupling of Gq protein with AT_1_-R mediate Ox-LDL/LOX-1-induced ERKs phosphorylation.

**Figure 7.**
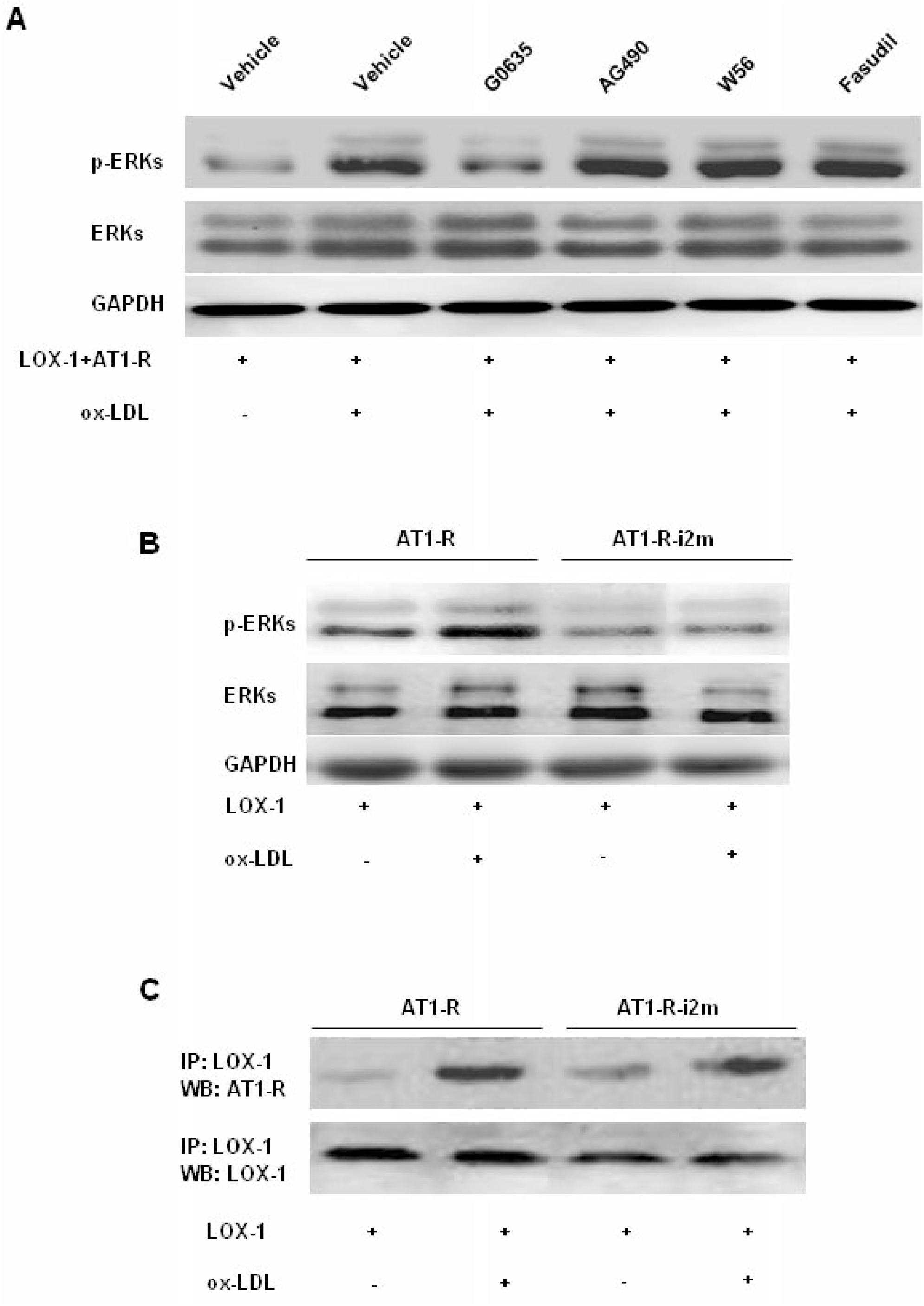
Effects of Gq protein on Ox-LDL-induced cellular events in LOX-1 and AT_1_-R-transfected cells. **A**, COS7 cells were transiently co-transfected with both *LOX-1* and *AT*_*1*_*-R* for 24 hrs. The Gq11 α subunit inhibitor G0635 (100 μg/L), Jak2 inhibitor AG490 (10 μmol/L), Rac1 inhibitor W56 (50 μmol/L), Rho kinase inhibitor Fasudil (10 μmolL) or vehicle was added to the cells. After 2 hrs, the cells were stimulated with Ox-LDL for 10 min. **B** and **C**, AT_1_-R or its mutant lacking ability of Gq protein coupling (AT_1_-Ri2m) were co-transfected with LOX-1 into COS7 cells and then the cells were stimulated by Ox-LDL. Phosphorylation of ERKs was detected by an anti phosphor-ERKs antibody. Total ERKs and GAPDH were used as loading controls. Membrane proteins were immunoprecipated (IP) with antibodies against LOX-1 and the immunecomplexes were subjected to Western blot (WB) analyses for AT_1_-R and LOX-1, respectively. Representative photograms from 5 independent experiments are shown.

### Ox-LDL induces direct binding of LOX-1 and AT_1_-R

To confirm whether LOX-1 and AT_1_-R bind directly or indirectly, we performed BiFC assay, which provides a novel approach for identification of the direct combination of two proteins modified by connecting to a specific protein construct in cultured cells [32, 33]. The following expression plasmids were used this assay: AT_1_-R connected at its C terminal to the subunit C (AT1/KGC) or the subunit N (AT_1_/KGN) of a GFP; LOX-1 connected at its N terminal to the subunit N or the subunit C of a GFP (LOX-1/KGN or LOX-1/KGC, respectively). Since green fluorescence can be detected only when the subunit N and the C of the GFP bound together, we co-transfected AT_1_/KGC with LOX-1/KGN or AT_1_/KGN with LOX-1/KGC into the COS7 cells and stimulated these cells with ox-LDL. After stimulation with Ox-LDL, green fluorescence was observed either in COS7 cells co-transfected by AT_1_/KGC and LOX-1/KGN (Figure 8A, middle) or in those with AT_1_/KGN and LOX-1/KGC co-transfection (Figure 8B, middle), indicating a binding of the subunit C or N of GFP tagged to the C-terminal of AT_1_-R with the N or C subunit of GFP connected to the N terminal of LOX-1, respectively. These results clearly suggest a direct binding of the C-terminal of AT_1_-R with the N-terminal of LOX-1 under the presence of Ox-LDL. However, AngII stimulation could not induce direct binding of AT_1_-R with LOX-1 (Figure 8A, B, right). It is usually believed that the receptors are expressed on the surface of cells. However, LOX-1 and AT_1_-R have been indicated existing also in cellular organelles [34, 35]. We here also observed expression and therefore interaction of the two receptors within the cells. To confirm whether the direct binding of LOX-1 to AT_1_-R functions, we also examined the phosphorylation of ERKs in the COS7 cells with the same transfections as above. Ox-LDL did induce the increases in the phosphorylation of ERKs in those cells co-transfected either with AT_1_/KGC and LOX-1/KGN or with AT_1_/KGN and LOX-1/KGC (Figure 8C). Interestingly, although AngII could not induce the direct association of AT_1_-R with LOX-1, it did activate ERKs in the similarly transfected cells (Figure 8D), further indicating that LOX-1 binding is not necessary for AngII-activated AT_1_-R and downstream signalling.

**Figure 8.**
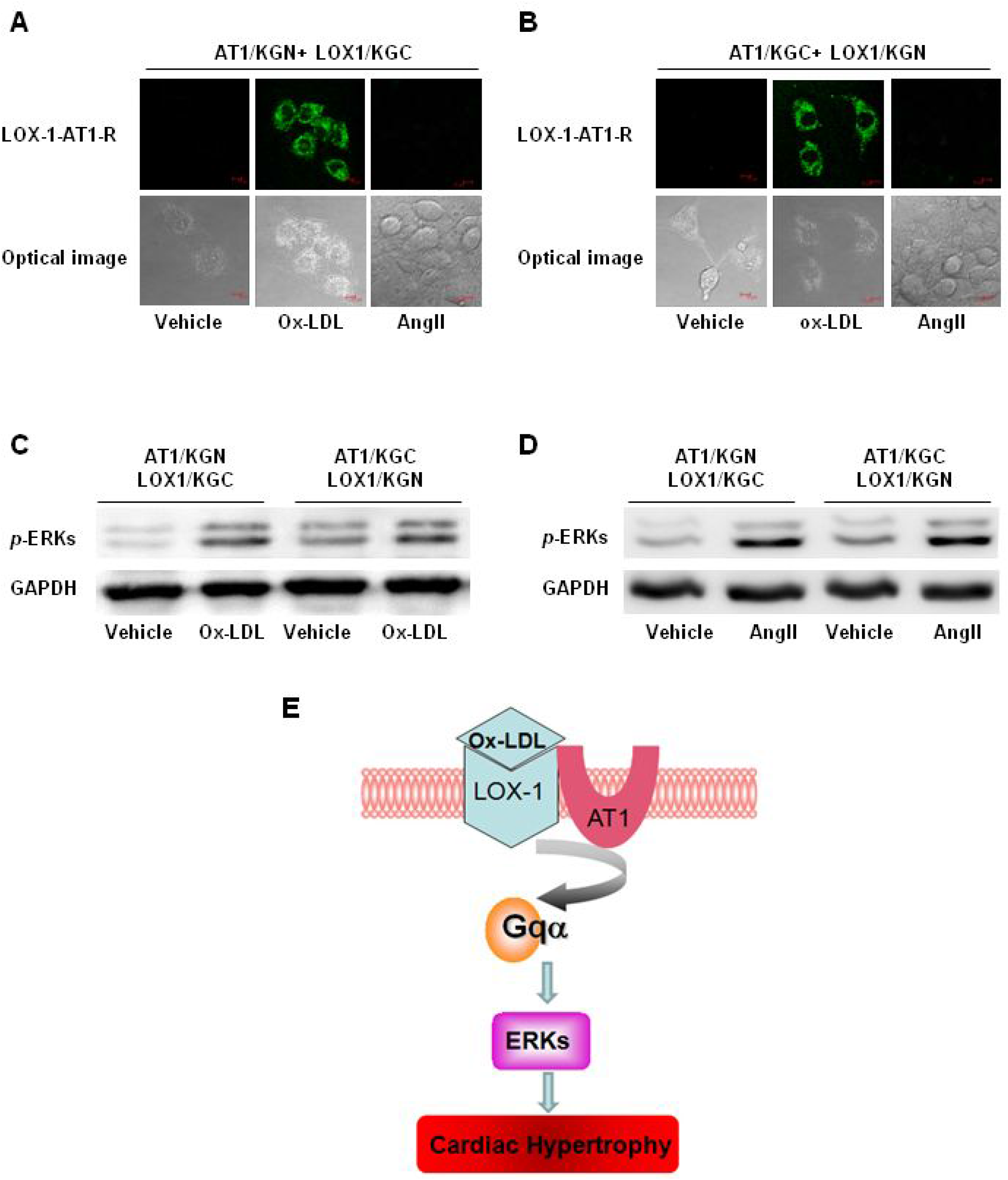
Ox-LDL induced-direct binding of LOX-1 and AT_1_-R. **A** and **B**, COS7 cells were co-transfected with AT_1_/KGC and LOX-1/KGN (**A**) or AT_1_/KGN and LOX-1/KGC (**B**), and stimulated by Ox-LDL, AngII or vehicle. Fluorescent signals indicate binding of the two receptors (top). Bottom, cell images through an optical microscope. Representative photographs from 5 independent experiments are shown. **C** and **D**, Ox-LDL- and AngII-induced ERKs phosphorylation. COS7 cells were co-transfected with AT_1_/KGC and LOX-1/KGN or AT_1_/KGN and LOX-1/KGC, and stimulated by Ox-LDL (**C**) and AngII (**D**) or vehicle (−) for 10 min. ERKs phosphorylation was detected by Western blotting using an anti phosphor-ERKs antibody. GAPDH was used as a loading control. Representative photograms from 5 independent experiments are shown. **E**. Proposed mechanism underlying ox-LDL-induced cardiac hypertrophic responses through binding of LOX-1 and AT1-R.

## Discussion

The present study provides two novel findings. Firstly, AT_1_-R could participate in Ox-LDL-induced cardiac hypertrophy without the involvement of AngII; Secondly, direct binding of LOX1 and AT_1_-R is one mechanism for Ox-LDL-induced hypertrophic responses.

It has been reported that Ox-LDL may be involved in cardiac injury including cardiac hypertrophy and heart failure [8–10]. It is also realized that Ox-LDL could induce a variety of cardiovascular injuries through LOX-1 [36]. In the present study, Ox-LDL induced both the *in vivo* and *in vitro* cardiomyocyte hypertrophy though LOX1, consisting with the previous concept. On the other hand, RAS including AngII and AT_1_-R have also been known to be involved in Ox-LDL/LOX-1-induced cardiovascular disorders [11, 28–30, 36]. Ox-LDL can induce increases in local AngII through LOX-1, thereby stimulates AT_1_-R. AngII induces oxidative stress and ROS generation via AT_1_-R transcription. Intense oxidant stress induces upregulation of NF-κB, leading to LOX-1 transcription and activation. The upregulation of LOX-1 further stimulates ROS generation which enhances AT_1_-R transcription and activation. However, in the present study, inhibition of either LOX-1 or AT_1_-R abrogated Ox-LDL-induced hypertrophic responses whereas reduction of AngII through inhibition of ACE only partially inhibited the responses stimulated by Ox-LDL, indicating that Ox-LDL can induce cardiac hypertrophy through LOX-1 and AT_1_-R rather than AngII. This idea is further supported by the results that Ox-LDL could induce cardiac hypertrophy even in AngII-deficient mice, which were also suppressed by LOX-1 and AT_1_-R inhibition. We therefore speculate here that Ox-LDL can induce cardiomyocyte hypertrophy through LOX-1 and AT_1_-R without the involvement of AngII.

Ox-LDL induced not only upregulation but also co-localization and association of LOX-1 and AT_1_-R, and inhibition of LOX-1 and AT_1_-R resulted in reduction of them each other, suggesting that Ox-LDL-induced cardiomyocyte hypertrophy is dependent on interaction of LOX-1 and AT_1_-R. Co-immuneprecipitation experiments potentially indicate that Ox-LDL-stimulation causes the association of LOX-1 with AT_1_-R even at the condition without AngII. Moreover, transfection of LOX-1 or AT_1_-R alone did not provide COS7 cells which lack endogenous LOX-1 and AT_1_-R with the ability to respond to Ox-LDL whereas overexpression of both of them did gave the cells this ability, further confirm the importance of LOX-1 and AT_1_-R interaction for Ox-LDL-induced cellular events. More importantly, our bimolecular fluorescence complementation assay clearly showed that LOX-1 and AT_1_-R could directly bind together, leading to activation of downstream molecules such as ERKs after stimulation with Ox-LDL. Previous reports indicated that there is an intimate relationship between the LOX-1 and the RAS: Ox-LDL increases ACE via LOX-1, resulting in an increase in AngII; AngII increases expression of LOX-1 through activation of AT_1_-R [11]. However, how Ox-LDL/LOX-1 interacts with AngII/AT_1_-R is still somewhat unclear. We here provide a potent evidence for the direct association of LOX-1 and AT_1_-R.

We have previously reported that besides AngII-dependent activation, AT_1_-R could be activated by mechanical stress independently of AngII and that the sensing sites for mechanical stretch within AT_1_-R are different from those for AngII stimulation [12, 20, 21]. In our present study, transfection of a mutant AT_1_-R (Q257A) lacking ability to sense mechanical stress abolished not only the Ox-LDL-induced hypertrophic responses but also the association of LOX-1 with AT_1_-R, indicating that Glu 257 is important for AT_1_-R to interact with LOX-1. K199Q mutant lacking ability of binding with AngII could not affect Ox-LDL-induced ERKs phosphorylation and the association of LOX-1 with AT_1_-R, consisting with the result showing that Ox-LDL-stimulated interaction of LOX-1 with AT_1_-R could be independent of AngII. However, Cys289Ala, another mutant lacking ability of response to mechnical stretch did not abolish both the Ox-LDL-induced ERKs phosphorylation and the receptor association, indicating that Cys 289 is not required for Ox-LDL-induced interaction of AT_1_-R with LOX-1. On the othe hand, the dominant negative mutant (K266A/K267A) of LOX-1 completely abrogated Ox-LDL-induced association of LOX-1 and AT_1_-R and hypertrophic responses in AT_1_-R-overexpressing cells, confirming that activation of LOX-1 is necessary to its association with AT_1_-R. We have previously reported that Gqα subunit and Jak2 mediate activation of AT_1_-R and ERKs by mechanical stretch [12]. Rac1 and RhoA have been known as the downstream molecules of LOX-1 [31]. In this study, inhibitor for Gqα subunit but not that for Jak2, Rac1 or RhoA inhibited Ox-LDL-stimulated ERKs phosphorylation in LOX-1/AT_1_-R-transfected COS7 cells, and transfection of the AT_1_-R mutant lacking coupling domain by Gq protein reduced Ox-LDL-stimulated ERKs phosphorylation but not association of AT_1_-R with LOX-1, suggesting that Gq protein exists as the downstream molecule of LOX-1/AT_1_-R interaction after Ox-LDL stimulation. Although we does not answer here why Gq but not Jak2 mediates ERKs activation from LOX-1/AT_1_-R, our another preliminary result show that Glu257 of AT_1_-R is important for the receptor to activate Gq protein but not Jak2 (Data not shown). Collectively, we speculate here that Glu257 of AT_1_-R and coupling of Gq protein with AT_1_-R account for the binding of LOX-1 to AT_1_-R and mediation of ERKs activation, respectively, after stimulation with Ox-LDL.

In summary, our present study shows another evince for AngII-independent Ox-LDL-induced activation of AT_1_-R mediated by LOX-1 binding. Although the precised molecular mechanism by which LOX-1 and AT_1_-R bind each other and lead to ERKs activation is not completely explored, our findings provide a new consideration not only for the study of G protein-coupled receptor activation but also for the clinical treatment of patients with hypercholesterolemia. It remains to be determined whether activation of AT_1_-R by Ox-LDL/LOX-1 without AngII occurs in other organs, and whether AT_1_-R blockers prevents organ damage more effectively than does ACE inhibitors in patients with both hypertension and hypercholesterolemia.

## Acknowledgement

We thank Jianguo Jia and Xia Sun at Fudan University for technical assistance. This work was supported by the grants from National Natural Science Foundation of China (81730009, 81941002), Laboratory Animal Science Foundation of Science and Technology Commission of Shanghai Municipality (201409005000).

## Author contributions

Y.Z. designed the study, L.L., N.Z., C.Y. and Y.Z. wrote the manuscript, Y.Z., C.Y. and J.G. originated the central idea and supervised the work. N.Z., L.K., J.W. and Q.W. performed *in vivo* experiments and analyzed the data, L.L., H.G., C.Y., G.Z. and Y.C. performed *in vitro* experiments and analyzed the data, H.A. and I.K. contributed the experiment materials supply, H.J., R.C., X.Y., A.S., Y.S., C.W., I.K. and J.R. analyzed the data and read the manuscript.

## Supplemental Materials

### Supplemental Methods

#### Echocardiography and blood pressure (BP) measurements

Transthoratic echocardiography was performed using 30 MHz high frequency scanhead (VisualSonics Vevo770, VisualSonics Inc. Toronto, Canada) [1–3]. All measurements were averaged for five consecutive cardiac cycles and were carried out by three experienced technicians who were unaware of the identities of the respective experimental groups. BP was evaluated as previously described [1–3].

Briefly, a micronanometer catheter (Millar 1.4F, SPR 835, Millar Instruments, Inc.) was inserted via the right common carotid artery into aorta. The transducer was connected to Power Laboratory system (AD Instruments, Castle Hill, Australia), and BP and HR were measured.

#### Morphology and histological analyses of hearts

Excised hearts were weighed, perfused with PBS followed by 4% polyformaldehyde for gross morphometry and fixed in 10% formalin for histological analysis. Paraffin embedded hearts were sectioned at 4 μm thickness, stained with hematoxylin and eosin (H-E). Cardiomyocytes were chosen from each section at a high magnification, and cross section area (CSA) of cardiomyocytes was measured by a video camera (Leica Qwin 3) attached to a micrometer in 20 different randomly chosen points from each cross section of LV free wall.

#### Lipoprotein preparation and modification of native LDL

Fresh human plasma, added citrate buffer solution containing 50 IU/ml heparin (pH 5.04), was subject to serial ultracentrifugation. After centrifugation, LDL was collected and dialyzed against phosphate buffered saline (PBS) buffer containing 200 μM EDTA for 48 hrs. Ox-LDL was prepared by incubation of LDL with 10 μM CuSO_4_ for 24 hrs at 37°C, followed by addition of 100 μM EDTA for 24 hrs, and then dialyzed in PBS buffer for 24 hrs at 4°C, changing the buffer every 8 hrs [4]. The concentration of protein was determined by the BCA protein assay kit. Oxidative modification of LDL by CuSO4 was assessed by the electrophoretic mobility of native and ox-LDL measured on 1% agarose gel [5]. The lipoprotein pattern was visualized by staining the gel with a lipid-specific stain.

#### Preparation and observation of aorta specimens

After 2 weeks of infusion with Ox-LDL, mice were sacrificed and the aorta was exposed. The fixed aorta was observed under the dissection microscope.

#### Measurement of plasma Ox-LDL

Blood was drawn from the retro-orbital venous plexus when the mice were killed. Ox-LDL was measured by using a mice Ox-LDL Elisa kit (CUSABIO BIOTECH CO., LTD., Japan) according to the manufacturer’s instructions.

#### Measurement of AngII concentration

Concentrations of AngII in serum of mice or culture medium of cardiomyocytes were measured using ELISA kits (Cusabio Biotech, Wuhan, China) according to the manufacturer’s instructions.

#### Preparation of Dil-labeled Ox-LDL and incubation with cultured cardiomyocytes

Ox-LDL was labeled with DiI as described previously [6]. In brief, the mixture of 75 μl DiI (3 mg/ml) added by 4 mg of Ox-LDL was incubated under sterile conditions at 37°C for 8 hrs. Labeled Ox-LDL was isolated by ultracentrifugation (100,000 g for 4 hrs). This procedure typically resulted in incorporation of 5-15 μg of DiI per mg of Ox-LDL. Cultured cardiomyocytes were incubated with DiI-Ox-LDL (50 μg/ml) for 24 hrs and the red fluorescence was detected under the confocal microscopy.

#### SiRNA of LOX-1 and AT1-R

SiRNA of *LOX-1* (5’-GCAUCUCAAGUUACAGUAATT-3’; 5’-UUACUGUAACUUGAGAUGCTT-3’), siRNA of *AT*_*1*_*-R* (5’-GCAAAGCUGUCUUACAUUATT-3’; 5’-UAAUGUAAGACAGCUUUGCTT-3’) and their respective scramble sequences were synthesized by PCR based method and purchases from Genepharma, Inc. (Shanghai, China). SiRNA was transfected into cultured cardiomyocytes at the optimized concentrations (0.66 μg/ml, ≈50% suppression) using the siPORT Amine Transfection Reagent (Ambion).

#### Western blot analyses

Total proteins were subjected to Western blot analysis for phosphorylation of ERKs, total ERKs, β-actin or GAPDH. The amounts of LOX-1 and AT_1_-R were examined after dividing the membrane fraction and the cytosolic fraction. Briefly, cells or tissue lysates were first centrifuged at 200 × *g* to remove nuclei. The supernatant was centrifuged at 15,000 × *g* for 30 min to pellet cell membrane. The total proteins or pelleted membranes were size-fractionated by SDS–PAGE and transferred to Immobilon-P membranes (Millipore). The blotted membranes were incubated with antibodies against phosphorylated ERKs (Cell Signaling Technology), total ERKs (Santa Cruz Biotechnology), LOX-1 (Abcam), AT_1_-R, β-Actin, GAPDH (Santa Cruz Biotechnology) and subjected to an ECL Detection system (GE healthcare).

#### Real-time RT-PCR

Total RNA was isolated from the LV tissues or cultured cells using TRIZol^®^ reagent according to the manufacturer’s instructions (Gibco BRL). After purification RNA was subjected to the real-time RT-PCR analysis on a BIO-RAD IQ5 multicolor detection system. Melting curves and quantization were analyzed using Light Cycler and Rel Quant software, respectively. A comparative CT method was used to determine relative quantification of RNA expression [7]. All PCR reactions were performed at least triplicate.

#### Fusion Proteins Design and BiFC Assay

The plasmids of AT_1_/KGC, AT_1_/KGN, LOX-1/KGN and LOX-1/KGC were constructed as previously described [8] using the CoraHue® FLUO-chase Kit (Medical&Biological Laboratories Co., LTD., Japan). Briefly, the cDNA of AT_1_-R and LOX-1 was amplified by PCR using pcDNA3-AT_1_-R and pcDNA3-LOX-1 as a template, respectively. The primers used are as follows: AT_1_-R: forward primer: 5’-AGCTCGGATCCACCATGTACCC-3’, reverse primer:5’-ACAAGCTTCTCCACCTCAGAACAAG-3; LOX-1: forward primer: 5’-CGGAATTCCGTGAATTTGGAAATGGCTTTTGAT-3’, reverse primer: 5’-CCGCTCGAGCGGTCACTGAGTTAGCAATAAATTTGCC-3’. The PCR products were digested with *EcoRI* and *xholI* and then inserted into vector of phmKGN-MC and phmKGC-MC to generate fusion plasmids. The following fusion plasmids were constructed: AT_1_/KGC, AT_1_-R tagged by subunit C of GFP at C terminals of the receptor; AT_1_/KGN, AT_1_-R tagged by subunit N of GFP at C terminals of the receptor; LOX-1/KGN, LOX-1 receptor tagged by subunit N of GFP at N terminals of the receptor; LOX-1/KGC, LOX-1 receptor tagged by subunit C of GFP at N terminals of the receptor. Since green fluorescence can be detected only when subunit N and C of GFP were bound together, AT_1_/KGC and LOX-1/KGN or AT_1_/KGN and LOX-1/KGC were co-transfected into the COS7 cells. Green fluorescence was detected under the confocal microscopy.

---

**Supplemental Figure 1.**
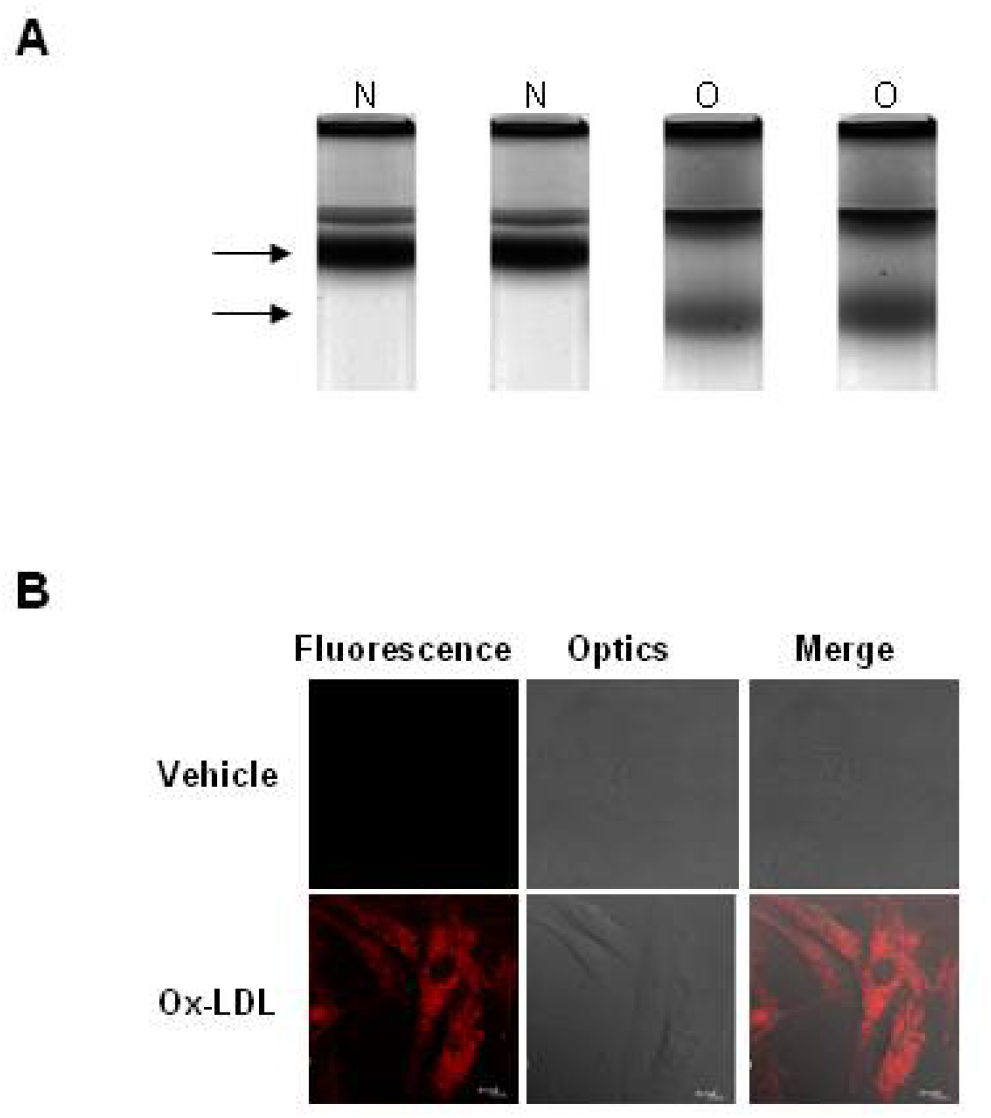
**A**, Electrophoretic mobility of native and Ox-LDL. The native (N) and the Ox-LDL (O) were electrophoretically analyzed on 1% agarose gel. The lipoprotein pattern was visualized by staining the gel with a lipid-specific stain. Representative photograms from 3 independent experiments are shown. The electrophoretic mobility defined as the ratio of the distances migrated from the origin was different between the modified LDL and native LDL. **B**, Uptake of Ox-LDL into cultured cardiomyocytes. Cultured neonatal cardiomyocytes of mice were incubated with vehicle (top) or Dil-labeled Ox-LDL (bottom) for 24 hrs. After fixation the cells were observed through a fluorescent (left) or an optical (middle) microscope. Right, merged images. Representative photomicrographs from 3 independent experiments are shown.

**Supplemental Figure 2.**
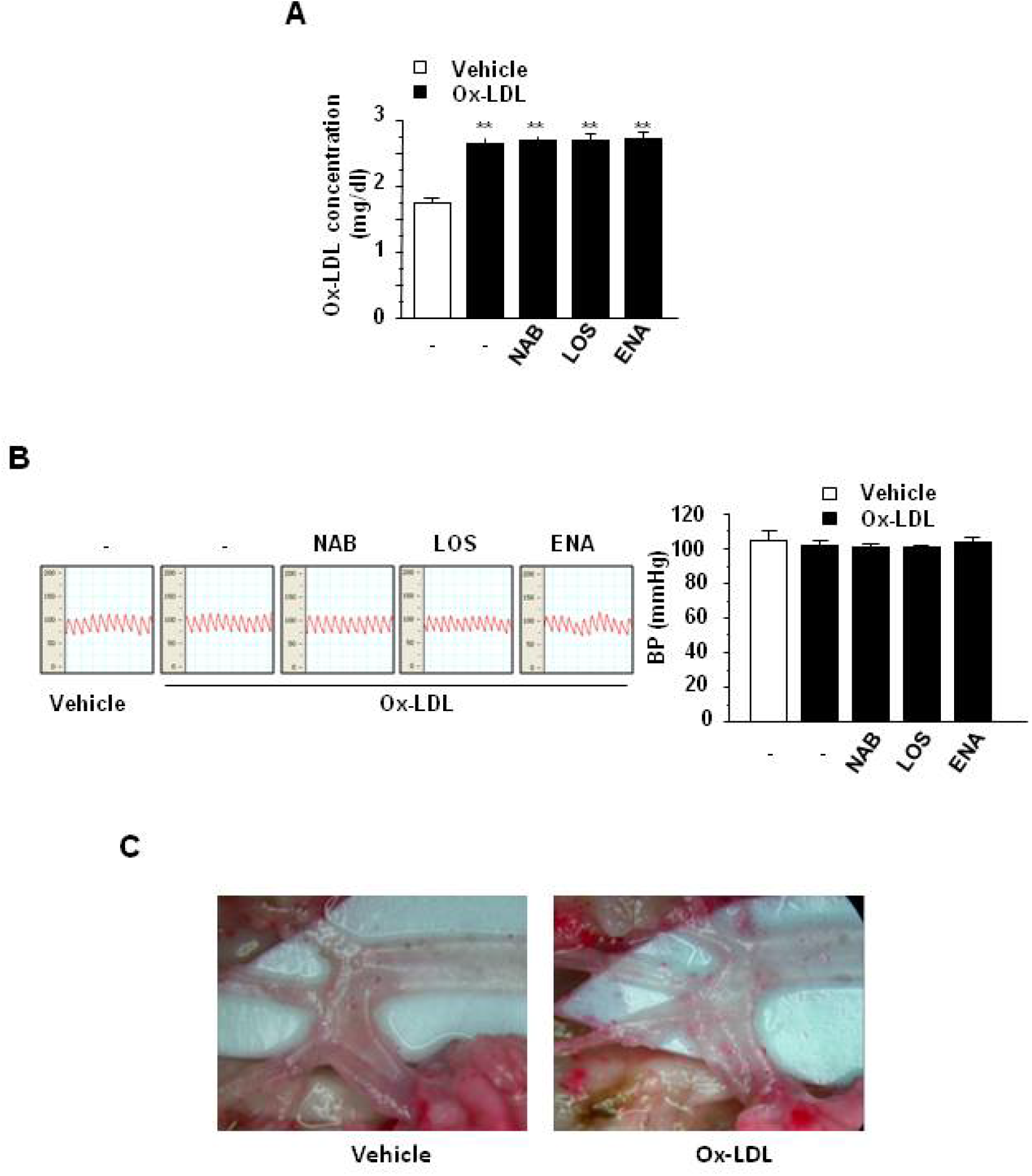
Adult male C57BL/6 mice pretreated with a neutralizing antibody of LOX-1 (NAB), Losartan (LOS), Enalapril (ENA) or vehicle (−), were infused with vehicle or Ox-LDL for 2 weeks. **A**, Measurement of plasma Ox-LDL. Plasma concentrations of Ox-LDL (mg/dl) were measured by Elisa method. All values are expressed as mean ± SEM of 5 mice (n=5). ** *p* < 0.01 vs. mice with vehicle. **B**, Blood pressure (BP). Representative recordings of BP from 5 mice are shown. All values are expressed as mean ± SEM of 5 mice (n=5). **C**, Representative photographs of gross appearance of aorta from 5 mice. Scale bar: 1 mm. There are no any atherosclerotic lesions observed in aorta.

**Supplemental Figure 3.**
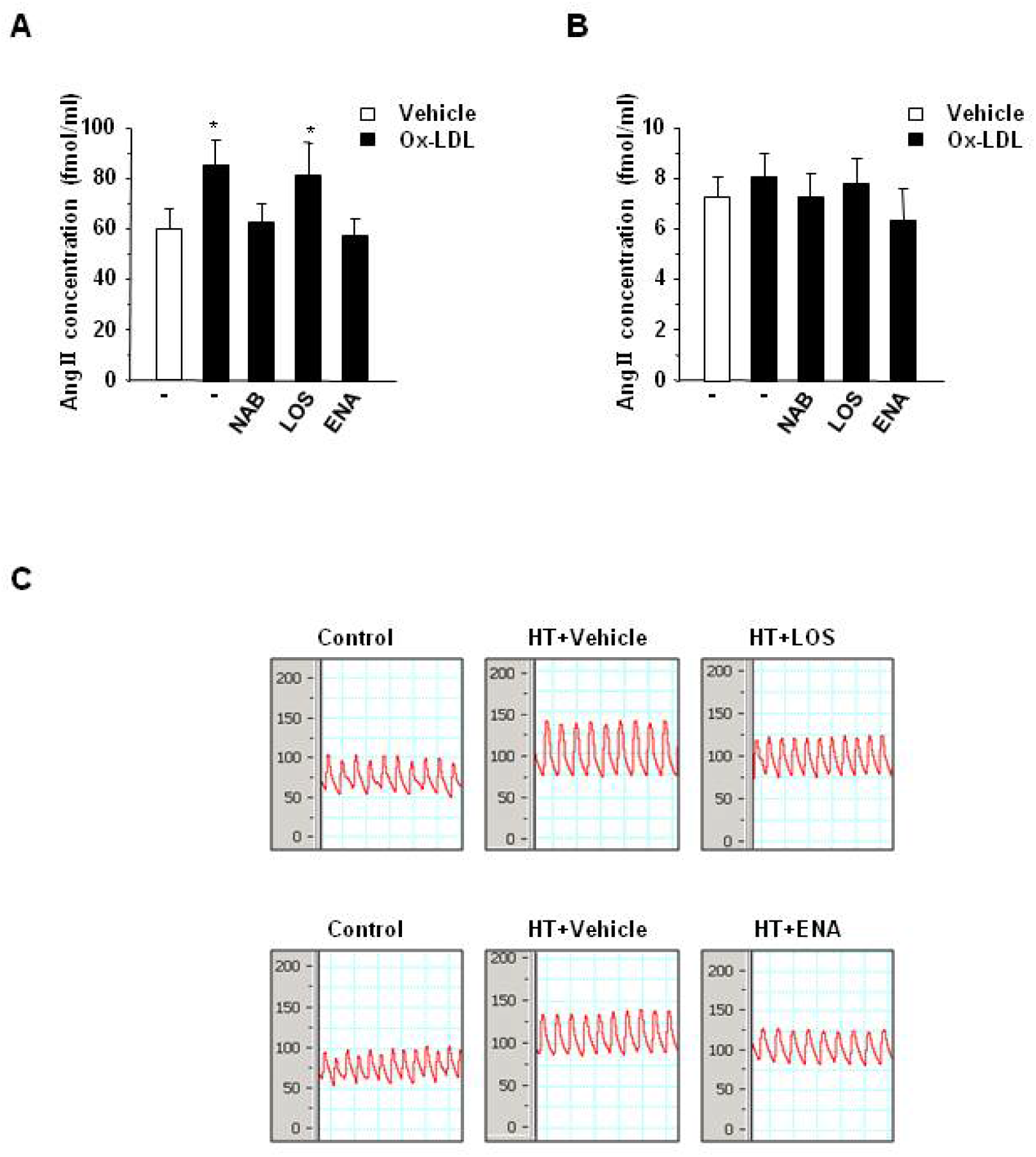
**A**, Adult male C57BL/6 mice pretreated with a neutralizing antibody of LOX-1 (NAB), Losartan (LOS), Enalapril (ENA) or vehicle (−), were infused with vehicle or Ox-LDL for 2 weeks. **B**, Cultured cardiomyocytes (CM) of neonatal mice, pretreated without (−) or with the NAB, LOS or ENA for 30 min, were exposed to vehicle or Ox-LDL for 24~48 hrs. AngII concentrations in plasma and culture medium were measured by Elisa method. * *p* < 0.05 vs. mice or cardiomyocytes with vehicle. **C**, Effects of Losartan and Enalapril on BP in hypertensive mice. Adult male C57BL/6 mice were subjected to an operation of constriction of abdominal aorta. Two weeks later, some of them were administered with saline (Vehicle), LOS or ENA for 2 weeks. Representative recordings of BP are shown. Control, mice without operation; HT, mice with hypertension after constriction of abdominal aorta.

**Supplemental Figure 4.**
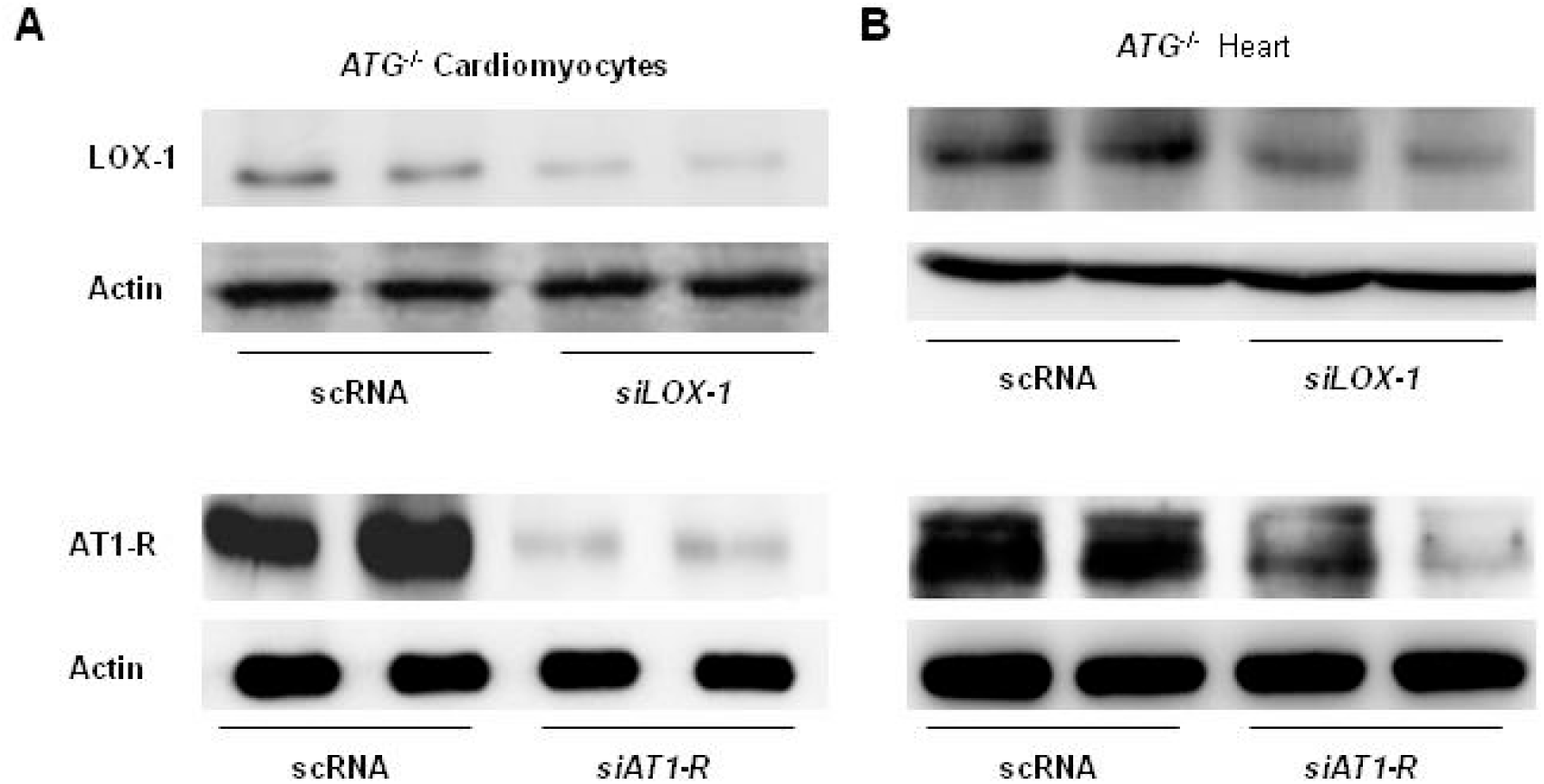
Inhibition of LOX-1 and AT_1_-R protein expressions by siRNA. **A**, Cultured cardiomyocytes of *ATG*^−/−^ mice were incubated with siRNA of LOX-1 (*siLOX-1*) (top) or AT_1_-R (*siAT*_*1*_*-R*) (bottom), or with respective scramble RNA (scRNA) (0.66 μg/ml) for 24 hrs. **B**, Adult male *ATG*^−/−^ mice were injected with AAV9-mediated shRNA of LOX-1 (*siLOX-1*) (top), AT_1_-R (*siAT*_*1*_*-R*) (bottom) or respective scramble RNA (scRNA). Two weeks later, mice were sacrificed and hearts were excised. The cellular membranes were isolated using the gradient centrifuge method and the protein expression of LOX-1 and AT_1_-R was detected by Western blotting. β-Actin in whole cellular lysate was used as a loading control. Representative photograms from 3 independent experiments are shown.

**Supplemental Figure 5.**
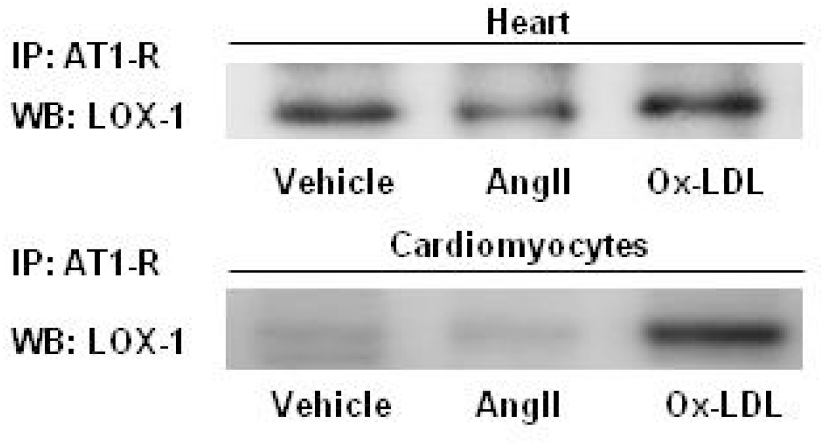
Different effects of AngII and Ox-LDL on LOX-1 and AT_1_-R association in heart and cardiomyocytes. Top, adult male C57BL/6 mice were infused with vehicle, AngII (200 ng/kg/min) or Ox-LDL for 2 weeks. Bottom, neonatal cardiomyocytes of mice were incubated with vehicle, AngII or Ox-LDL for 24 hrs. Membrane proteins extracted from LV of mice or cardiomyocytes were immunoprecipated (IP) with antibodies against AT_1_-R and the immunecomplexes were subjected to Western blot (WB) analyses for LOX-1. Representative photograms from 5 mice or 5 independent experiments are shown.

**Supplemental Figure 6.**
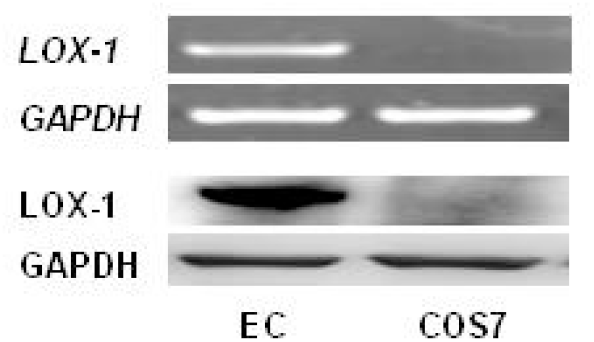
Examination of LOX-1 expression in COS7 cells. mRNA was isolated from cultured COS7 cells and vascular endothelial cells (EC) and the expression of *LOX-1* was examined by RT-PCR (top). The cellular membranes were isolated from cultured COS7 cell and EC using the gradient centrifuge method and the protein expression of LOX-1 was detected by Western blotting (bottom). GAPDH was used as the loading controls. Representative photograms from 3 independent experiments are shown.

**Supplemental Figure 7.**
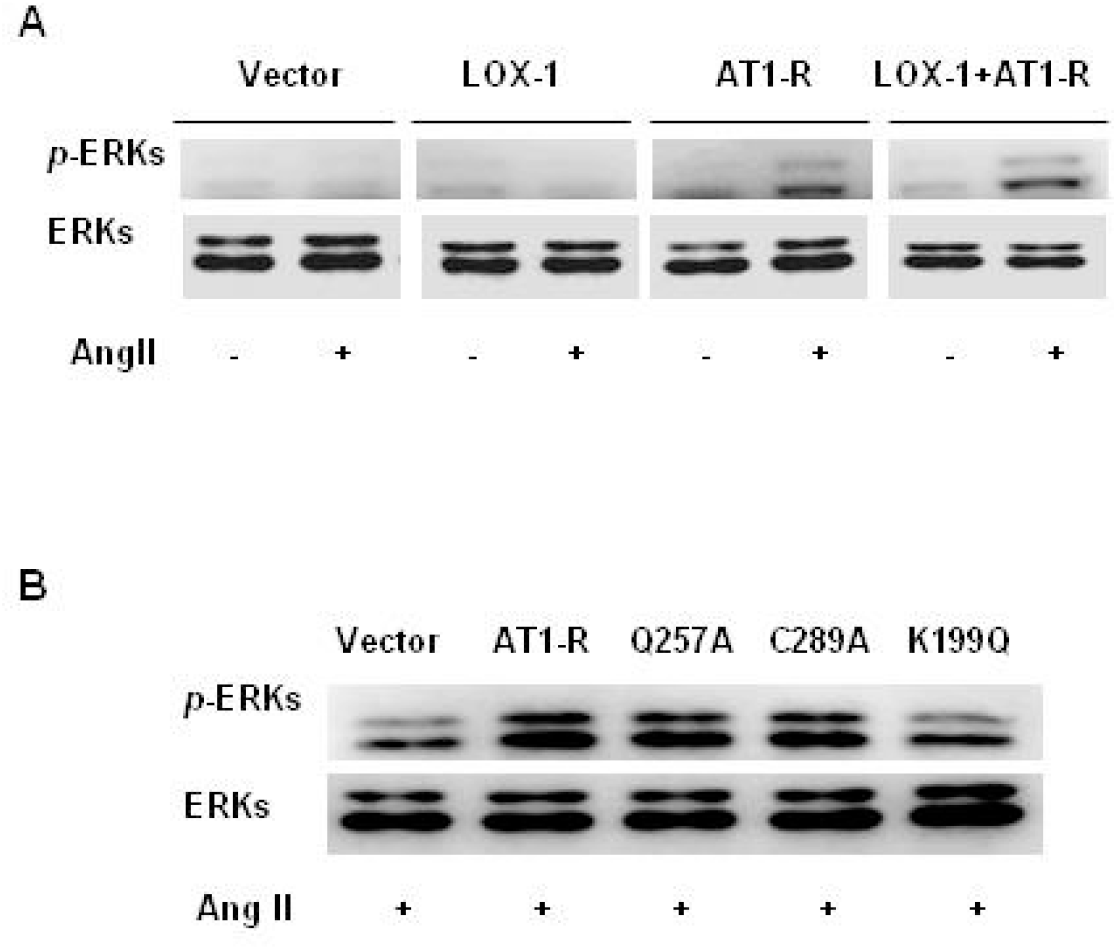
Effects of co-transfection of LOX-1, AT_1_-R or their mutants on ERKs phosphorylation. **A**, COS7 cells were transiently transfected with empty vector, *LOX-1* or *AT*_*1*_*-R* alone, or co-transfected with both *LOX-1* and *AT*_*1*_*-R* for 24 hrs and stimulated by AngII or vehicle for 10 min. **B**, COS7 cells were transiently co-transfected by *LOX-1* with *AT*_*1*_*-R* mutants K199Q, G257A or C289A for 24 hrs and stimulated by AngII for 10 min. Phosphorylation of ERKs was detected by an anti phosphor-ERKs antibody. Total ERKs were used as a loading control. Representative photograms from 5 independent experiments are shown.

